# Uncovering the modified immunopeptidome reveals insights into principles of PTM-driven antigenicity

**DOI:** 10.1101/2021.04.10.438991

**Authors:** Assaf Kacen, Aaron Javitt, Matthias P. Kramer, David Morgenstern, Tomer Tsaban, Adam Solomon, Guo Ci Teo, Felipe da Veiga Leprevost, Eilon Barnea, Fengchao Yu, Arie Admon, Lea Eisenbach, Gal Cafri, Ora Schueler-Furman, Yishai Levin, Alexey I. Nesvizhskii, Yifat Merbl

**Affiliations:** Department of Immunology, Weizmann Institute of Science, Rehovot, Israel; De Botton Institute for Protein Profiling, Nancy and Stephen Grand Israel National Center for Personalized Medicine, Weizmann Institute of Science, Rehovot, Israel; Department of Pathology, University of Michigan, Ann Arbor, Michigan, USA; Department of Microbiology and Molecular Genetics, Institute for Medical Research Israel-Canada, Faculty of Medicine, The Hebrew University, Jerusalem, Israel.; Faculty of Biology, Technion - Israel Institute of Technology, Haifa 32000, Israel; Department of Computational Medicine and Bioinformatics, University of Michigan, Ann Arbor, Michigan, USA; Sheba Medical Center, Ramat Gan, Israel

## Abstract

Antigen processing and presentation are critical for modulating tumor-host interactions. While post-translational modifications (PTMs) can alter the binding and recognition of antigens, their identification remains challenging. Here we uncover the role PTMs may play in antigen presentation and recognition in human cancers by profiling 29 different PTM combinations in immunopeptidomics data from multiple clinical samples and cell lines. We established and validated an antigen discovery pipeline and showed that newly identified modified antigens from a murine cancer model are cancer-specific and can elicit T cell killing. Systematic analysis of PTMs across multiple cohorts defined new haplotype preferences and binding motifs in association with specific PTM types. By expanding the antigenic landscape with modifications, we uncover disease-specific targets, including thousands of novel cancer-specific antigens and reveal insight into PTM-driven antigenicity. Collectively, our findings highlight an immunomodulatory role for modified peptides presented on HLA I, which may have broad implications for T-cell mediated therapies in cancer and beyond.

**Significance:** Major efforts are underway to identify cancer-specific antigens for personalized immunotherapy. Here, we enrich the immunopeptidome landscape by uncovering thousands of novel putative antigens that are post-translationally modified. We define unique preferences for PTM-driven alterations affecting HLA binding and TCR recognition, which in turn alter tumor-immune interactions.

**Conflict of interest statement:** Authors declare no conflicts of interest.

## Introduction

Targeting tumor antigens that are bound to Major Histocompatibility Complex (MHC) molecules holds great promise for T cell therapies and immunotherapies. Peptides derived from foreign pathogens and self-proteins, which have undergone disease-related changes, such as mutations^1–5^, may elicit an immune response. Similarly, post-translational modifications (PTMs) such as phosphorylation, citrullination, or glycosylation^6–12^ may also occur on presented antigens, and have been reported to modulate antigen presentation and recognition^13^. For example, such changes in the antigenic landscape were reported in clinical phospho-proteomic analysis of breast and lung cancer, uncovering differential activation of cellular pathways^14,15^. However, with more than 200 different types of PTMs, and the technical difficulties in detecting them, whether and to what extent such PTM-driven alterations expand our landscape of antigenic targets in cancer, remained under-explored.

Current approaches for neoantigen discovery rely mostly on genomic or transcriptomic data^16^, combined with computational prediction tools for Human Leukocyte Antigen Class I (HLA I) binding17–21. Such approaches are geared towards identifying neo-antigens generated by mutations or non-canonical amino acid sequences. Since they are focused on the pre-translational level, they lack information on the state of modification of the peptides. Another approach relies on the identification of HLA I-bound peptides by immunoprecipitation of the MHC/ HLA-peptide complex from the surface of cells and eluting the bound peptides prior to mass spectrometry (MS)-based analysis (i.e. immunopeptidomics). MS analysis and the identification of peptides is done by comparison of the peptides detected by the instrument to a reference dataset containing all the possible theoretical peptides across the proteome. Thus, to detect PTMs on such peptide, requires the relevant reference sequence that contains the same mass shift imparted by the modification. As each additional modification increases the number of theoretical peptide possibilities in the search space exponentially, the search time becomes a limiting factor. Many approaches to cope with the exponential growth of the search space when searching for PTMs have recently been implemented (open search^22^, denovo^23^, ‘), in various mass spectrometry analysis tools (MetaMorpheus^24^, PEAKS PTM) ^23–26^. To date, however, the vast majority of PTMs, and combinations thereof, have not been examined in the immunopeptidome.

To address these challenges and examine the potential landscape of modified peptides that are bound to HLA I in a systematic and unbiased manner, we developed a PROtein Modification Integrated Search Engine (PROMISE). Our computational pipeline allows for combinatorial detection of multiple PTMs without prior biochemical enrichment. To test PROMISE we analyzed the modified immunopeptidome landscape in a murine model of cancer. We could confirm our predictions by peptide spectra validation followed by killing assays towards cancer-associated modified peptides, *ex vivo* and *in vivo*. By examining data generated from 210 samples, including patient-derived tumor samples and cancer cell lines, we found thousands of novel modified HLA I-bound peptides which generated cancer -specific signatures. Notably, some of these modified peptides reside within known cancer-associated antigens or cancer driver genes, offering a novel class of antigens which may be further examined in the context of immunotherapy. By systematically analyzing the locations of PTMs on MHC-eluted peptides we uncovered PTM-driven motifs across many haplotypes, in many cases altering the anchoring positions or the middle region of the peptide, which is associated with T cell recognition region. We further confirmed these observations by using structural 3D modeling. Finally, by extending our analysis to a breast cancer cohort from the Clinical Proteomic Tumor Analysis Consortium (CPTAC^14^) database, we revealed cancer- and site-specific modifications. Such sites were identical to the ones we found in modified antigens, bringing insight into metabolic changes and the altered modification landscape induced by the cancerous state. Collectively, our systematic identification of modified peptides and their impact on HLA I binding and recognition should broaden our understanding of the effects PTMs may have on defining tumor-host interactions.

## Results

### Establishment of the Protein Modification Integrated Search Engine (PROMISE)

The assignment of peptides that were detected by the MS instrument to their cognate amino acid sequence is based on the reference proteome that is provided to the analysis software. As such, peptides that are eluted from HLA I may only be identified if they match a specific ‘theoretical’ peptide in a defined search space (i.e. reference proteome). Peptides that are detected by the MS but cannot be matched or assigned to any sequence are considered as the ‘dark matter of the proteome’ ^27^. The dark matter of the proteome may include all sequences that deviate from the encoded amino acid sequence of proteins, such as mutations, non-canonical translation products, fusion proteins, spliced peptides or PTMs ^28–32^. For the latter, identifying modified peptides that may be presented on HLA I, in a systematic manner, remains a challenge due to the huge search space of endogenous peptides with the numerous possibilities of protein modifications. In recent years, several approaches were implemented to cope with this challenge (MetaMorpheus^24^, PEAKS PTM), offering state-of-the-art solutions to peptide assignment. Here, we developed PROMISE (PROtein Modification Integrated Search Engine) which relies on the ultrafast MSFragger^33^software for comparison between the theoretical peptides and the peptide captured in the instrument (see supplementary pipeline documentation for details). PROMISE simultaneously searches HLA immunopeptidomics data against multiple modification types that are not identified by standard analysis (Figure 1A). Modifications identified by PROMISE can indicate either PTMs that represent the altered protein state in the cell or modifications that may have been introduced during sample processing (e.g. carbamidomethylation ^34^and deamidation^35,36^). Nevertheless, incorporation of diverse types of modifications to the search space allows us to choose the best match for the detected peptides and assign peptides that would otherwise not be identified. Only peptides that match better to a theoretical peptide with a modification than all other possible matches to the encoded amino acid sequences in the proteome (see materials and methods), are defined as modified peptides by PROMISE. To identify a broad range of PTMs, we defined 29 modification combinations of 12 modification types (36 mass shifts; Supplementary data 1) on 16 different amino acids and protein termini (termed hereafter ‘multi-modification search’). These include modifications such as methylation, acetylation, phosphorylation, citrullination, ubiquitination, sumoylation, oxidation, deamidation, cysteinylation and carbamidomethylation.

**Figure 1.**
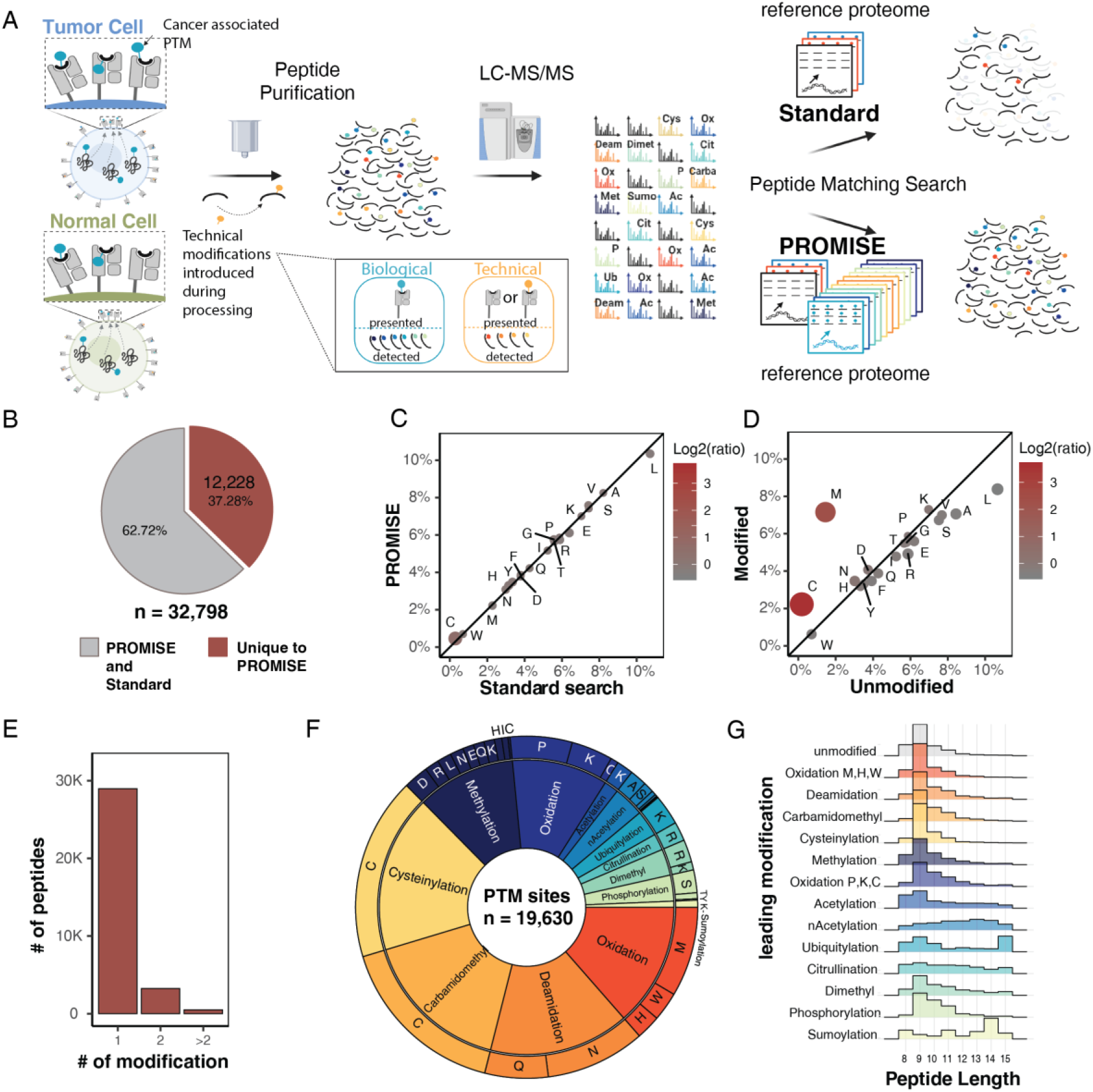
Computational pipeline for global search of PTMs on HLA I-bound peptides enriches identifications. **(A)** Protein Modification Integrated Search Engine (PROMISE) allows for the systematic detection of modifications on HLA I-bound peptides (**B**) Pie chart of modified peptides identified in the standard and multi-modification search performed on multiple immunopeptidomics datasets. Out of 32,798 modified peptides identified in the analysis, 37.29% were unique to PROMISE. (red). **(C + D)** The amino acid composition of peptides identified was compared for the standard and PROMISE search (C) or the unmodified and modified subsets of peptides in the PROMISE search (D). Circle size and color indicate the log2 transformed ratio of amino acid abundance between the two subsets. (**E**) Peptides identified in PROMISE are binned by number and type of modification. (**F**) When viewed by modification site, 19,630 positions were uniquely identified by PROMISE in the immunopeptidomics datasets analyzed. These sites are then presented in a pie chart divided by modification type, and amino acid modified. (**G**) Peptide length distribution (density as a percentage of total peptides) per modification type.

### Global search of PTMs on Human sample of HLA I-bound peptides enriches identifications

We next sought to utilize PROMISE to identify putative modified antigens in the context of human cancers. To that end, we ran multi-modification search to analyze previously published high-resolution HLA I immunopeptidomics data (PRIDE identifications: PXD004894^7^, PXD000394^37^, PXD006939^38^, PXD003790^39^, PXD009738^40^) of patient tumors tissues (n = 35) or healthy adjacent tissue (n = 5), cancer cell lines (n = 13), and TILs (n = 2). To identify peptides for which the modified state was a better match to the spectrum, we compared our results to the original search criteria, which in most of our datasets included methionine oxidation and protein N-terminus acetylation (termed hereafter ‘standard search’). In both cases, we used a subgroup FDR at 5% by splitting spectra into three different groups based on modification state (see methods), ensuring we are not increasing identifications merely by altering the false positive rate. The multi-modification search identified 32,798 modified peptides (Supplementary data 1). In total, 12,228 of the modified peptides identified were unique to the multi-modification search, thereby enriching the pool of immunopeptides identified (Figure 1B). While the amino acid composition of the immunopeptidome was similar between the standard search and PROMISE, we saw an enrichment in amino acids that can carry modifications when comparing the modified and unmodified peptide subsets (Figure 1C,D). For example, as previously described^41^, cysteines are consistently under-represented in immunopeptidomics analyses, yet constitute 2% of the modified immunopeptidome (Figure 1D). As expected, most of the modified peptides carried only one modification (Figure 1E). In total, we identified 19,630 modification sites (from 12,228 peptides) that were unique to PROMISE, 88% of which contain modification types that are not included in a standard search (Figure 1F). We next analyzed the length distribution per modification type and observed that acetylation, citrullination, dimethylation, sumoylation, and ubiquitylation are longer on average than the unmodified subset (Figure 1G).

### An unbiased search of 29 modifications highlights PTM-driven preferences

Given our global view of post-translationally modified peptides, we wished to explore if a given PTM has the tendency to be in certain positions within the peptide. To capture the motifs of the full modified peptide repertoire, we used a global FDR correction (Supplementary data 3). A broad view across different types of modifications reveals that some modifications have a distinct site preference (Figure 2A). For example, as previously shown^6,7^, serine phosphorylation predominantly falls in the 4^th^ position of the peptide while methylation is distributed evenly across the peptide (Figure 2A, light green). Further, we found that oxidation and cysteinylation are enriched at the end of the peptide (towards the c-terminus), and carbamidomethyl is enriched in the third position, while cysteinylation is under-represented at the second position.

**Figure 2.**
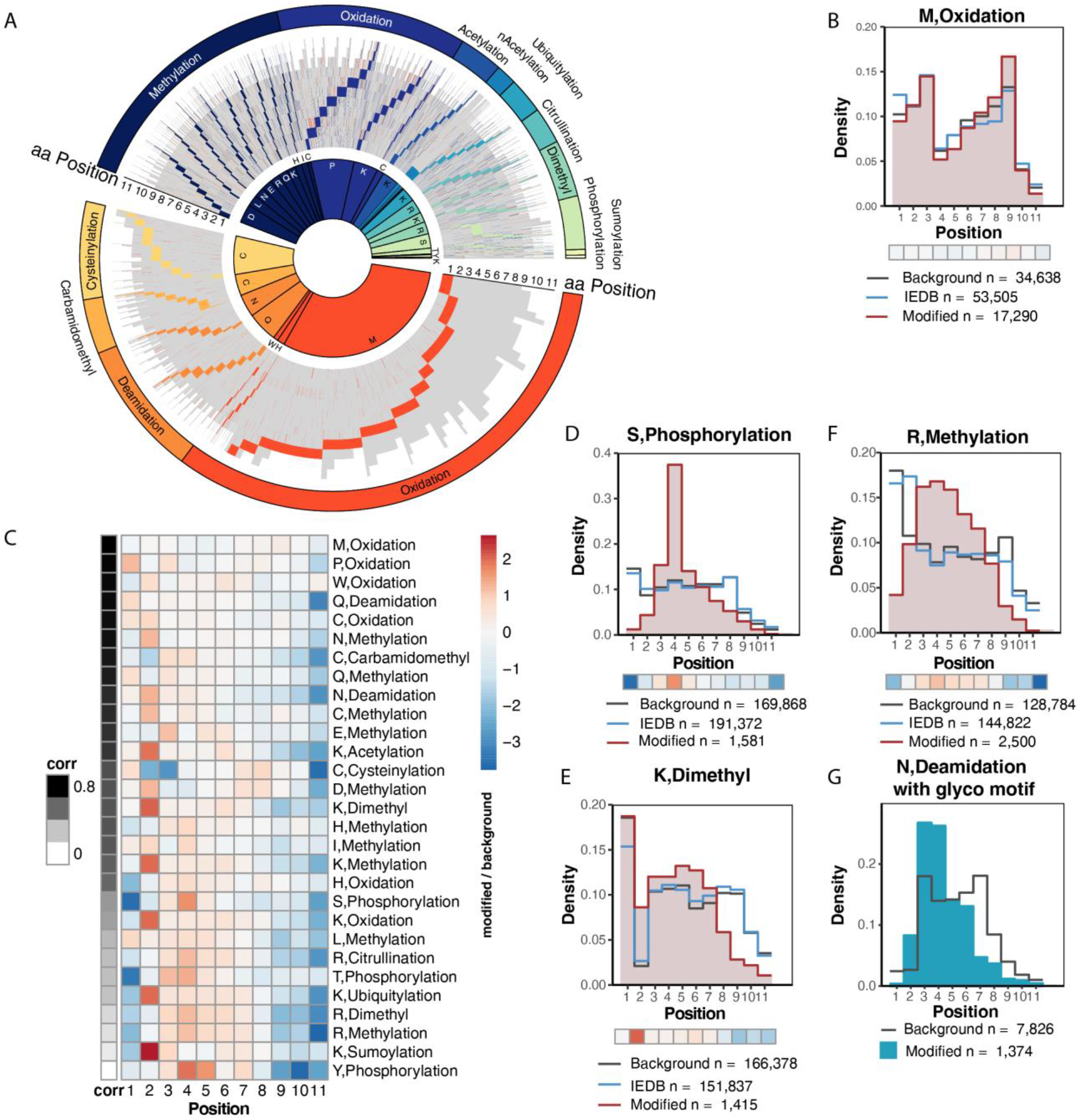
PTM-driven binding preference highlighted through unbiased search of 29 modifications. (**A**)All the peptides with modifications identified in the reanalysis of the Bassani et al37 dataset by PROMISE (n = 12,268 peptides) are sorted by the modification type and position in the peptide. Each line represents a distinct peptide in grey with the modification site(s) colored. (**B**) Correlation between oxidized methionine position distribution and the un-modified methionine distribution is very high (Pearson 0.96, p value 1.05e-6), and as expected from a technical artifact the distributions are not significantly different (F-test; p value = = 0.3543). **(C)** Modification distributions are sorted by the correlation between the modified amino acid and the un-modified background. A low correlation means the PTM distribution is distinct from the unmodified background, suggesting a PTM-driven motif. (**D-F)** We compared the modified amino acid position distribution (“Modified”, red) to the distribution of the unmodified amino acid that carries this modification in the analyzed datasets (“background”, grey) or identified in the IEDB 42 database (“IEDB”, blue). Major differences between those distributions suggest that the modified amino acid has position preferences not solely determined by the properties of the unmodified amino acid. Below each histogram, the fold change between the modified AA and unmodified AA distribution is presented as a heatmap bar (red indicates overrepresentation of the modified AA relative to the unmodified distribution). (D) Distribution of serine shows the phosphorylated form falls predominantly falls in the 4th position and significantly different from the unmodified serine distribution (F-test; p value < 2.2e-16). (E) Lysine residues are underrepresented at the second position of the peptide, however the distribution of the dimethylated form is enriched at the second position compared to the background (F-test; p value < 2.2e-16). (F) Methylated arginine is enriched in positions 3 to 7 compared to background arginine (F-test; p value < 2.2e-16). **(G)** Deamidated asparagine with a glycosylation motif is enriched in position 3 and 4 compared to background asparagine (F-test; p value < 2.2e-16).

Next, we explored whether the distribution of these PTMs is distinct from the underlying distributions of the amino acid residues that they modify. In addition, we also examined an unbiased and broader background distribution by collectively defining all of the reported epitopes in the IEDB^42^ database. As expected, when examining a modification which is widely generated by sample processing/handling, like methionine oxidation, the correlation between the oxidized methionine position distribution and the un-modified methionine distribution is very high (Pearson 0.96, p value = 1.05e-6) (Figure 2B). This suggests that the modification occurred randomly across the peptide during sample preparation (F-test; p value = 0.3543). As this was not the case for all the PTMs, we ordered all of the PTMs we detected based on the correlation of their distribution to the background (Figure 2C). This metric highlights PTM-motifs which may alter the HLA binding preference or TCR recognition. Peptide binding to HLA I molecules depends on the biochemical properties of both the peptide and HLA I structure. The most critical residues for HLA I binding are the ones that fit into the anchor pockets in the HLA I groove, typically the second and carboxy-terminal positions^43^. By contrast, T-Cell receptors recognition motif is determined by the HLA I-peptide complex and therefore most strongly influenced by the residues in position 3 to 7 of the HLA I-bound peptide^44,45^. In the presented matrix for example, known motifs, such as the tendency of serine phosphorylation modification at position 4^6,7^, were also emphasized as low correlation in this analysis (Pearson 0.41, p value = 0.21) as there was a strong deviation between the phosphorylation and underlying serine distributions (Figure 2D; F-test; p value < 2.2e-16). This was identified despite any experimental or computational enrichments for specific modifications, as we used a broad search that was not modification-specific. Beyond confirming known motifs, we also identified novel ones. For example, lysine residues are generally underrepresented in the HLA I binding pocket at the second position of the peptide. However, modified lysine residue distributions (e.g. acetylated and methylated lysine) do not produce the same pattern (Figure 2E). This suggests that unmodified lysine residues in the anchoring position are unfavorable for HLA I binding and that the modified state of a lysine residue may be preferred. In contrast, modified arginine such as di/methylated arginine and citrullination are over-represented in positions 3 to 7, and therefore may impact the T-cell receptor recognition^44^ (Figure 2F), as was previously shown for other modifications types. Interestingly, while cysteine modifications on peptides in MS analyses are considered to be introduced by sample processing, in our analysis of the HLA I landscape they have a distinct distribution motif where cysteine carbamidomethyl is enriched in positions 3-4 and cysteinylation is enriched in positions 7-8 (Figure 2C).

The deamidation of asparagine residues occurs naturally at glycosylated sites on proteins^36^ and these sites have a strong consensus sequence motif of asn-x-ser/thr. Peptides with N-deamidation and the glycosylation motif, suggesting they are biological in origin, show a distinct tendency to be located on the 3rd and 4rd position of the peptides (Figure 2G; F-test; p value < 2.2e-16). Another interesting group of peptides are those modified with ubiquitin (UB) or ubiquitin-like (UBL) tails (Figure 2A). These are likely a result of the ubiquitin undergoing degradation with its substrate, leaving several residues as a remnant modification^46^. With the aid of PROMISE, we were able to identify eight different remnant mass shifts of different lengths derived from at least three different UB/UBL types across multiple datasets (Supp. Fig 1). While these patterns emerged by globally examining the modification landscape, it is clear such frequencies may be affected by the distribution of HLA haplotypes on which they are presented.

### HLA I binding properties are altered by the modification state of the presented peptide

The biochemical binding properties of specific HLA haplotypes are the strongest determinants of peptide motifs. To examine whether the PTM-driven motifs we have detected is associated with specific haplotypes, we re-analyzed mono-allelic HLA immunopeptidomics data from Abelin et al^18^ (MassIVE: MSV000080527). We conducted the same multi-modification search as described above (Supplementary data 1) on the spectra obtained in this study. Indeed, we could identify unique motifs that were haplotype-dependent, using the unmodified amino acid distribution as a background. To focus on the most prominent features, we defined a ‘site score’ such that enrichment in the anchor positions will result in a positive score while enrichment in the middle of the peptide will result in a negative score. In case the PTM is present in many positions in the peptide, the score will be close to zero and we cannot classify the tendency of the modification to be in a specific area. We then clustered the biological PTMs and haplotypes contained in the dataset by their site score (Figure 3A). This analysis revealed that the same PTM might affect peptide-MHC-TCR interactions differently for different haplotypes. Intriguingly, among the specific HLA haplotypes that we analyzed, we found several HLA associations with human diseases. For example, HLA A*0301 was linked to increased risk for multiple sclerosis ^47^ and HLA B*5101 was linked to Behçet’s disease^48^. Our analysis identified both haplotypes to be highly enriched with PTMs in the region that is predicted to affect TCR recognition. HLA-A0201 was previously reported to show a protective effect in EBV-related Hodgkin lymphoma patients^49^ and in our analysis was enriched with modifications on the anchoring position of the peptide. While it remains to be examined whether certain PTMs play a role in disease-associated manifestations, it has been reported that low HLA binding of disease-associated epitopes can be increased by PTMs^50^.

**Figure 3.**
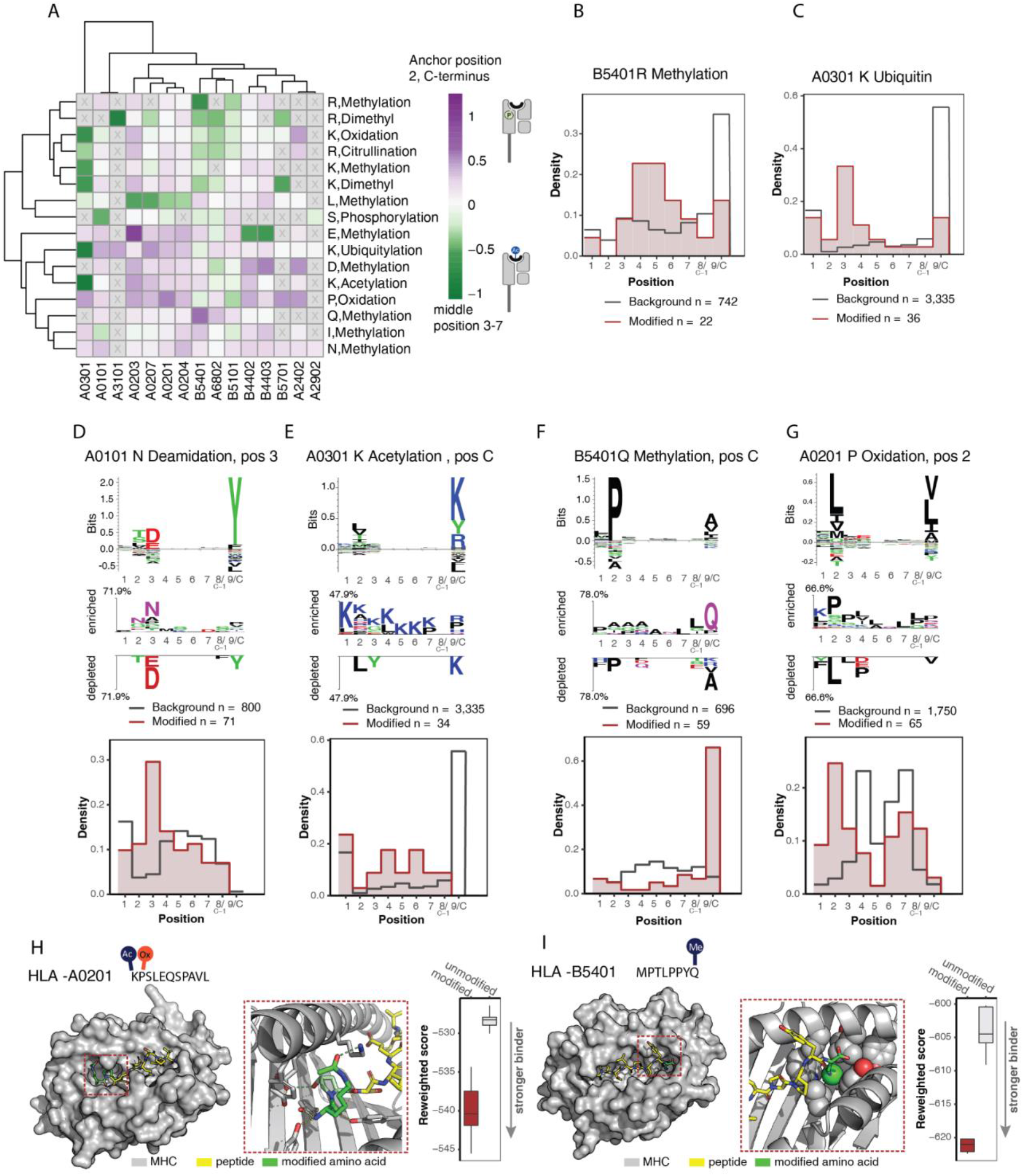
PTM-driven HLA motifs. (**A**)A recognition area score was calculated (see methods) to determine the tendency of a given modification to be located in the MHC anchor position (purple) or center of the peptide (green) for a given HLA haplotype. (**B-C**) The histogram then represents the modified amino acid frequency in each position (red) compared to the unmodified amino acid background (grey). C-terminus and C-1 are presented at position 8 and 9 (see methods). (B) Methylated arginine in haplotype B5401 is enriched in position 4-6. (C) Ubiquitin tail on lysine is enriched in position 3 of haplotype A0301. (**D-G**) Motif of the reported unmodified epitopes in the IEDB database for the indicated haplotype (top). The canonical modified motif was then compared to the amino acid motif for a given modification (middle). The histogram then represents the modified amino acid frequency in each position (red) compared to the unmodified amino acid background (grey). C-terminus and C-1 are presented at position 8 and 9 (see methods).(D) *Chemical mimics motif:* Aspartic acid is favored in the A0101 binding motif at position 3. (E) *Binding interference:* acetylated lysine is under-represented in the C-terminus of haplotype A0301. (F, G) *novel motif:* methylated glutamine at the peptide C-terminus in haplotype B5401 and oxidized proline at the anchor position 2 of haplotype A0201 creates favorable binder peptides. (**H + I**) Rosetta FlexPepDock structural models of the interactions between the modified peptide (yellow sticks) including the modified amino acid (green)and the MHC molecule (grey surface \ cartoon). The effect of the modified amino acid is shown in detail in the zoom-in picture. FlexPepDock reweighted score was calculated for the interaction between the MHC and modified or unmodified peptide. More negative score indicates a more stable interaction. (H) Interaction between K(ac)P(ox)SLEQSPAVL and haplotype HLA-A0201,hydrogen bonds introduced by the modification shown as dashed green lines. Other hydrogen bonds between peptide and receptor are shown in yellow dashed lines. (I) Interaction between MPTLPPYQ(me) and haplotype HLA-B5401: The glutamine methyl group is shown as green sphere, MHC interacting residues shown as gray spheres. The modified peptide shows significant lower predicted affinity (measured as FlexPepDock reweighted score).

PTM enrichment in the middle of the peptide, potentially affecting TCR recognition, could be observed with methylated arginine on haplotype B5401 (Figure 3B) or ubiquitin tail on lysine on haplotype A0301 (Figure 3C). PTM enrichment in an anchor position were classified into three groups: The first group is comprised of c*hemical mimics*, where the modified amino acid is biochemically similar to a different amino acid that was known to be part of the motif. For example, we identified an enrichment of deamidated asparagine in position 3 of the haplotype A0101 motif. Deamidated asparagine is chemically similar to aspartic acid which appears in the A0101 binding motif at position 3 (Figure 3D). As we could not find an unmodified peptide carrying asparagine bound to this haplotype, this result suggests that the modification occurred on the peptide before being bound to the HLA, possibly due to the removal of a glycosylation^51^, and the modified asparagine enables the binding of the peptide to the HLA. Enrichment of deamidated asparagine and glutamine at HLA haplotype A6802, B4402, and B4403 (Supplementary data 4) are additional examples of *chemical mimics*. The second group contains PTMs that cause *binding interference*. This group is defined by PTMs that sterically hinder the interaction of the peptide with the MHC haplotype, creating an unfavorable binder. For example, acetylated lysine is under-represented in the C-terminus of haplotype A0301 (Figure 3E) compared to the unmodified background. Importantly, we found this observation to hold true for all of the modified lysines detected in this haplotype, suggesting that the modification of the carboxy-termini could be an immune evasion mechanism. Other examples for *binding interference* are methylated glutamic acid at anchor position 2 of haplotype B4402/3, and dimethylated arginine at the C-terminus position of haplotype A3101 (Supplementary data 4). The third group is *novel motifs* where the modified amino acid creates a favorable binder peptide that is different from the known unmodified motif. It was shown that phosphoserine can replace glutamic acid at anchor position 2 of haplotype B4002^9^. In our dataset, we detect methylated glutamine at the peptide C-terminus in haplotype B5401 (Figure 3F) and oxidized proline was observed at the anchor position two of haplotype A0201 (Figure 3G). The latter observation is common to the whole haplotype superfamily A02 (Supplementary data 4).

Next, we evaluated the possibility of a novel PTM binding motif using structural modeling. To that end, we chose two representative modified epitopes identified as binders of haplotype A0201 and one representative epitope identified as a binder to haplotype B5401. All of them are shared across cancer cell lines and patient’s tumor samples. We used Rosetta FlexPepDock^52^ to model the structure of the interactions of these novels MHC-binding PTM motifs, K(ac)P(ox)SLEQSPAVL, KP(ox)LKVIFV and MPTLPPYQ(me). For each such motif, we modeled both the modified and unmodified peptides and compared their calculated binding energies and structures (“Reweighted score”). In all cases, the interactions between the MHC and the modified peptide interactions were predicted to be considerably stronger, suggesting the complex is more stable than the non-modified counterpart (Figure 3H, I, Supp. Fig 2) in agreement with the predictions from PROMISE immunopeptidomics analysis. In the case of peptide K(ac)P(ox)SLEQSPAVL binding to HLA-A*0201, our model suggests that the hydroxyl group of peptide P(ox)-2 forms a stabilizing hydrogen bond with receptor E-87 (Figure 3H). Overall, our models recapitulate an interaction similar to a solved structure of HLA-A2 in which T-2 forms hydrogen bonds with receptor K-90 and E-87 (1TVB^53^). As for K(ac)-1, in some of our models it interacts with the aliphatic part of receptor K-90, while in others it further stabilizes the peptide. In the case of peptide MPTLPPYQ(me) binding to HLA-5401, Q-8 is positioned in the highly hydrophobic pocket that binds the canonical aliphatic c-terminal peptide position. Methylation allows the otherwise polar (negative) side chain of glutamine to approach (“fill”) the pocket and thereby stabilize the complex (Figure 3I). Together our findings show that modified peptides are distinct from their coded counterparts in haplotype preference, binding motifs and structural interactions. We therefore wished to examine whether PTMs may also alter antigen reactivity.

### Modified peptides identified by PROMISE are specific to cancer and can elicit CD8 T cell reactivity

To test whether PROMISE-identified peptides may be antigenic, we performed immunopeptidomics on MC38 murine colon cancer cells. PROMISE multi-modification search analysis revealed 2,803 peptides, 36% of them with at least one PTM (Figure 4A). To determine which of the modified peptides are unique to the MC38 cancer cell line and are not detected in healthy tissue we utilized PROMISE to analyze immunopeptidomics data of healthy mouse tissues, from Schuster, H. et al^54^, using the multi-modification search (Figure 4B). We chose modified peptides that did not appear in healthy tissues and were not reported in IEDB (Supplementary data 5). We then proceeded to synthesize and validate these peptides. All the synthesized peptides were confirmed to match the original identification through manual annotation and scoring of spectrum similarity (Figure 4C, Supp. Fig 3). These included peptides with N-terminal acetylation, citrullination, dimethylation, methylation, phosphorylation and SUMOylation remnants (GGT) (Figure 4D). We chose from the validated MHC-bound peptide identified by PROMISE 3 modified peptides: TALFHTD(me)D(me)I, SYR(Dimethyl)GNYGGGGGG and SR(Me)GGGD(Me)QGYGSGR (from Protein Tyrosine Phosphatase Non-Receptor Type 14, PTPN14 DExH-Box Helicase 9, DHX9 and RNA Binding Motif Protein 3, RBM3). To examine whether the modified peptides are antigenic, we performed two independent assays to show (I) peptide-specific *in vivo* killing, and (II) the tumor killing potential of peptide-specific lymphocytes.

**Figure 4.**
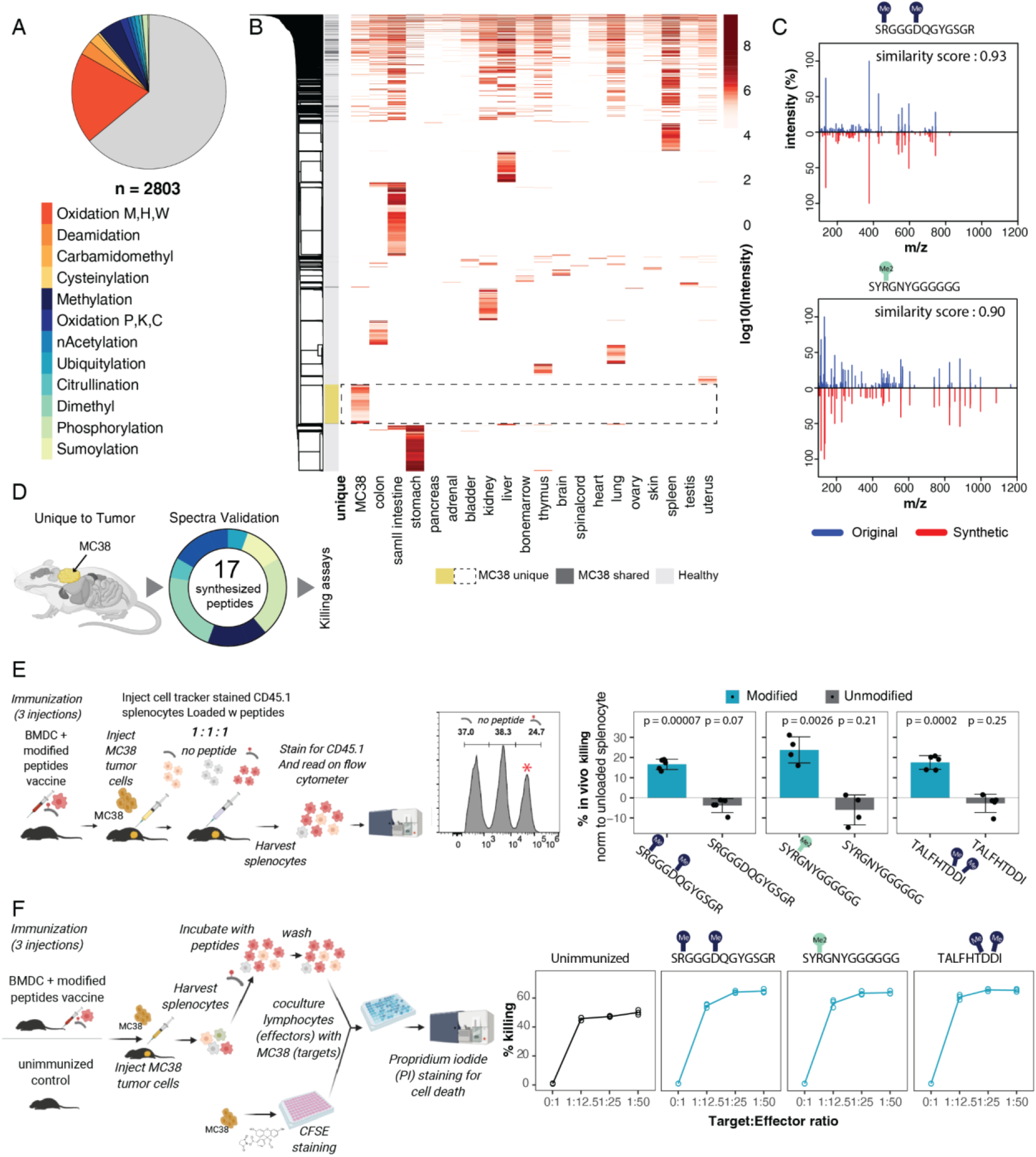
Modifications on MC38 associated peptides modulate MHC binding and alter CD8 T cell activation and killing. **(A)** PROMISE analysis of MC38 immunopeptidome identified 2,803 peptides. **(B)** Modified peptides from PROMISE multi-modification search analysis of 19 different tissues from healthy mice (Schuster, H. et al dataset) as well as MC38 cancer cell line (total number of peptides = 8,547) are presented by their MS intensity (color code in red) in the relevant column specifying the tissue of origin in which they were identified. Tissues are clustered by similarity revealing shared modified peptides between MC38 and healthy tissue (dark gray panel on the left) and unique peptides to MC38 (yellow on the left panel; dashed square). **(C)** Examples of modified peptides identified in the datasets, each peptide was synthesized (Peptide 2.0 Inc) and its spectrum was capture using mass spectrometry. A similarity score was calculated between the synthesized spectrum (red) and the original spectrum in the dataset (blue). **(D)** Peptide selection overview: 17 peptides were identified in the MC38 cancer line and not in the healthy mouse tissues and passed spectra validation. These peptides were then screened for MHC binding and reactivity and several peptides were assessed for their tumor killing potential. The pie charts show the types of modifications and the number of peptides which were tested**. (E)** In vivo Killing. Splenocytes from CD45.1 mice were pulsed with either modified or unmodified peptides, or unpulsed as control, and differentially labeled by specific dyes. Labeled splenocytes were injected in 1:1:1 ratios of modified peptide pulsed: unmodified peptide pulsed: unpulsed into MC38-tumor bearing mice that had been immunized prior to tumor injection with either modified or unmodified peptides. 18 hours later, the differentially labeled CD45.1 splenocytes were harvested and counted. The killing percentage of splenocytes that were peptide-pulsed is calculated relative to the killing of unpulsed splenocytes (n = 5 mice). Negative values reflect less killing than unpulsed control, and killing by splenocytes pulsed wit unmodified peptides did not differ significantly from killing of unpulsed cells (p values from one sample student’s t test, mu = 0). **(F)** Ex vivo killing. Splenocytes were harvested from MC38 tumor bearing mice that had been immunized with the pooled three modified peptides, and then sensitized ex-vivo with the indicated specific peptides for 5 days. For comparison, splenocytes were harvested from unimmunized tumor-bearing mice and were not pulsed by peptides. Then, the splenocytes were cultured with target MC38 cells and the percentage of MC38 target cell killing was determined. light blue: immunized; black: unimmunized. (n=3).

To test the *in vivo* reactivity against the three selected modified peptides, we performed BMDC-mediated immunization, followed by *in vivo* killing assay,adapted from ^55,56^ with the following addition: following the immunization, the mice were injected with MC38 tumor cells and allowed to grow for 18 days. Then, we used splenocytes loaded with either the modified peptide (labeled with high concentration of cell tracker dye), unmodified peptide (labeled with low concentration) or unloaded splenocytes (labeled with a medium concentration) and injected them in 1:1:1 ratio into immunized mice. After 18 hours, specific killing was measured (Supp. Fig. 4A). The three modified peptides induced between 20-30% specific killing (Figure 4E), suggesting that these peptides can induce peptide-specific cytotoxicity *in vivo*. By contrast, the percentage of specific killing did not significantly differ between the splenocytes loaded with unmodified peptides and the unloaded splenocytes control, indicating that the modification itself impacted the response. Finally, to demonstrate that the identified epitopes induce tumor-cell killing, we examined the ability of peptide sensitization to elicit target cell killing. Specifically, we immunized mice with the peptides and injected MC38 cells to form tumors. We then harvested splenocytes from immunized-tumor bearing mice and sensitized them ex-vivo by incubating with each of the specific peptides for 5 days. For comparison, we harvested splenocytes from MC38 tumor bearing mice which were not immunized and these splenocytes were left unsensitized. Then, the splenic lymphocytes (immunized and sensitized or control) were co-cultured with MC38 tumor cells to assess cytotoxicity. Notably, we found that immunization and sensitization with any of the modified peptides induced a 10%–30% increase in tumor cell killing compared to killing elicited by lymphocytes from tumor bearing mice which were not sensitized (Figure 4F). This serves as a proof-of-concept that modified antigens may elicit specific T cell reactivity.

### The modified HLA I landscape uncovers hundreds of tumor-associated modified antigens

Given the growing interest in identifying antigenic targets for immunotherapy, we examined whether we identified modified peptides originating from cancer-associated or testis antigens. We identified 98 peptides that originated from a protein annotated as a testis antigen (CT Antigens Database^57^, Figure 5A - left). For these, we examined their mRNA expression in TCGA data of the matching cancer types (Supp. Fig 5A) and found a subset to be overexpressed in the tumor tissue when compared to the adjacent controls (Supp. Fig 5B). We also identified 300 peptides that are highly shared between patients and across cancer cohorts (Figure 5A, right). Many of these proteins are also annotated as oncogenes, cancer drivers, or tumor suppressors^58^, highlighting the importance of studying the state of these proteins in tumor immunogenicity. None of these cancer-associated target peptides would have been identified without including PTMs in the protein search space.

**Figure 5.**
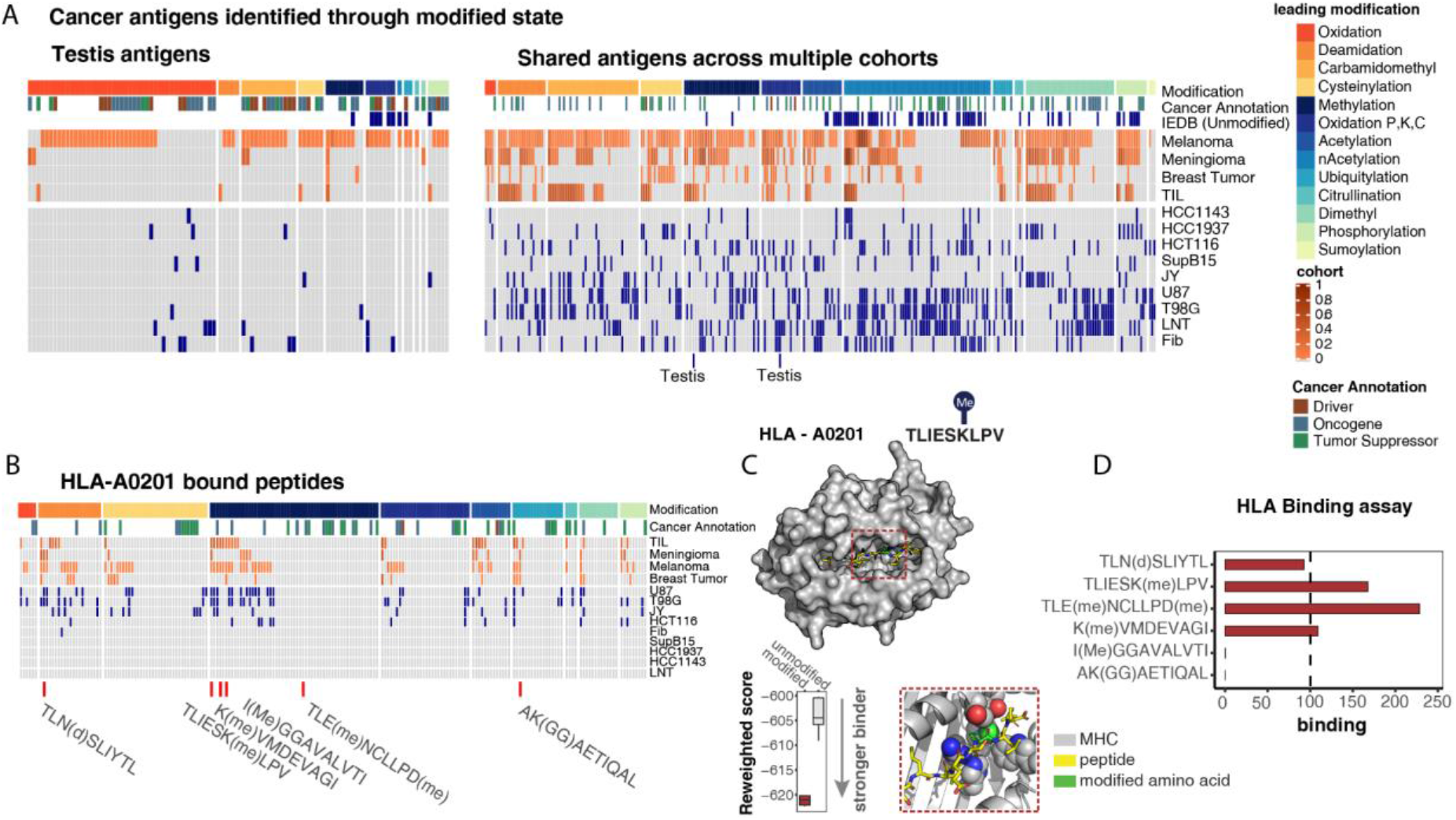
Tumor-associated antigens and cancer-specific classes of PTMs. (**A-B**) Each list of antigens is sorted by the modification of the peptide. For each peptide we mark the cancer annotation (driver, oncogene, tumor suppressor) as documented in CancerMine 58 if the peptide was reported in IEDB 42 in its unmodified state, and if it is a cancer-testis antigens. For a cohort of patient samples (orange) the color indicates the percentage of the patients the peptide was identified in. For cancer cell lines (blue) the color indicates that the peptide was detected. (A) Modified cancer-testis antigens list (n= 98, left) and a list of shared antigens (n=300, right) identified through the modified state.(B) A list of HLA-A0201 bound modified peptides that were not reported in the IEDB database. Rosetta FlexPepDock structural model of the interactions between TLIESK(me)LPV (yellow sticks) and the HLA-A0201 molecule (grey surface / cartoon). The methylated lysine (green) is packed against hydrophobic residues of the MHC molecule (gray spheres). The modification created a more stable interaction with the MHC molecule. (**D**) Six modified peptides from the list in B’ were tested for binding affinity through ProImmune *in vitro* binding assay.

To focus on the modified peptides we identified with PROMISE asscociated with a specific haplotype I, we filtered for modified peptides that were identified in immunopeptidomics from a HLA-A0201 cell line and that were not identified in IEDB in their unmodified form (Figure 5B). We then examined whether the difference in the detection of the modified peptides and their unmodified counterparts was due to their relative ability to bind HLA-A0201. Using structural modeling, we were able to show that the methylation on the lysine in position 6 of TLIESKLPV is located between 3 other positively charged residues (H-98, R121, and H-138; Figure 5C). Methylation of K-6 removes its positive charge and thereby alleviates electrostatic repulsion. In addition, the methyl group is nicely packed into the hydrophobic MHC groove. This then causes a more stable peptide-MHC interaction as reflected in a lower reweighted score. To assess the role of peptide modification in altering MHC binding we synthesized 6 peptides and examined binding using a binding assay (ProImmune, see methods). Of the peptides synthesized, 4 modified peptides were confirmed as HLA I binders (Figure 5D).

### Cancer-induced alterations in metabolic and PTM states are presented in the antigenic landscape

To determine whether these signatures are also specific to the cancer state in clinical settings, we analyzed immunopeptidomics data from a cohort of Triple-Negative Breast Cancer (TNBC) and adjacent tissue ^40^. Within this cohort, we found 2,771 modified peptides. We assessed whether there are classes of PTMs that are more frequent in the immunopeptidome of the tumor samples versus their adjacent controls. We found several modifications that were significantly reduced in frequency in the tumor immunopeptidome, including carbamidomethyl and citrullination (Figure 6A). Further, we found that cysteinylated peptides are significantly increased in the tumor immunopeptidome. The tumor and adjacent tissues were processed and analyzed together and therefore are not expected to have differential effects in modifications that were generating merely by the processing procedures. As such, the results likely signify changes in modifications elicited by the biological system. These changes may reflect alterations in metabolic pathways or peptide processing. For example, it is known that TNBC is addicted to cysteine^59,60^, potentially explaining the increase in cysteinylated immunopeptides.

**Figure 6.**
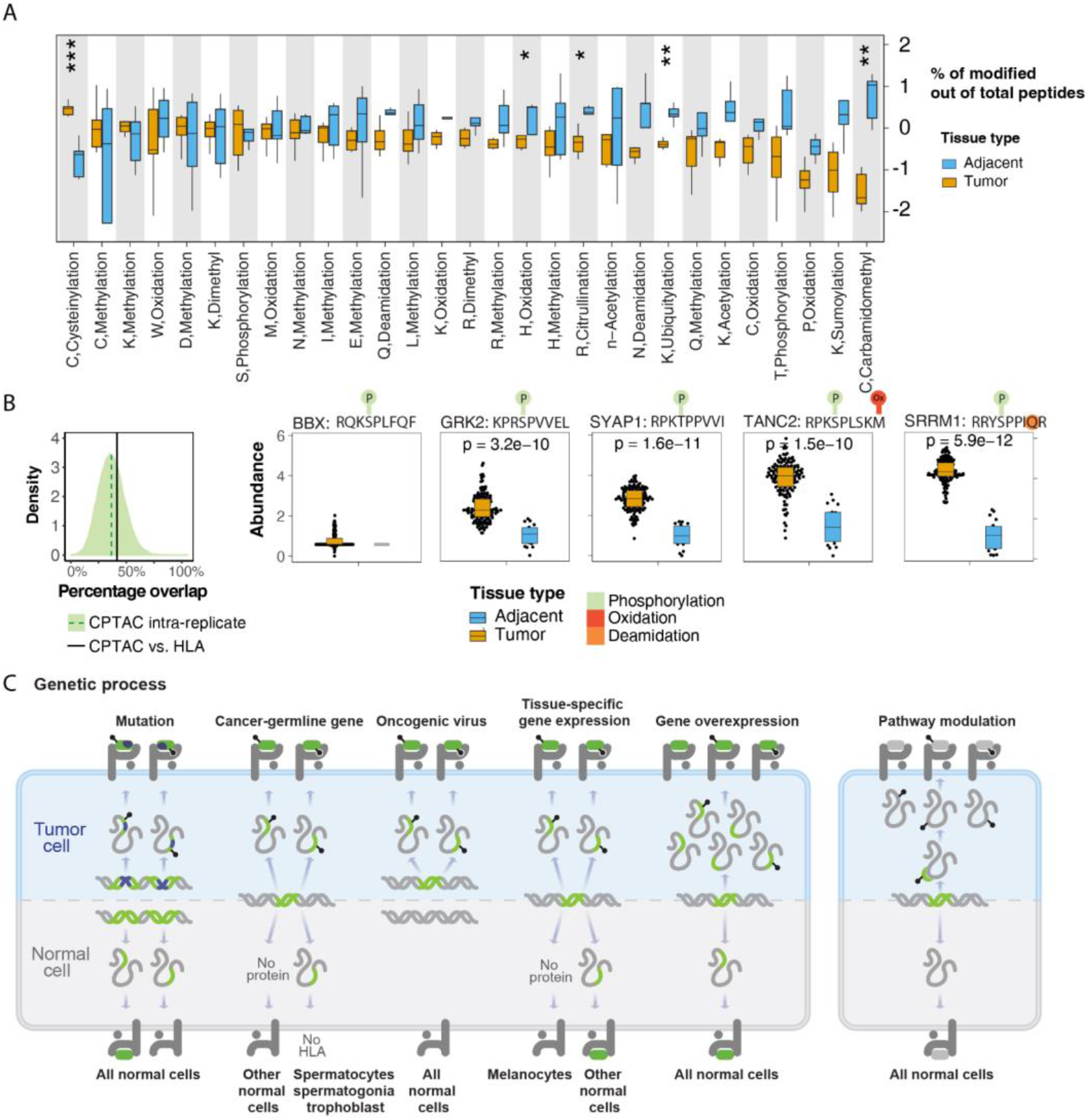
Modified HLA-bound peptides create cancer-specific signatures. (**A**)The percent of immunopeptides identified with each of the indicated modifications was calculated for a cohort of triple-negative breast cancer tumors and adjacent tissue (Ternette, N. et al40). The modifications are sorted from the most enriched in the tumor tissue at the left to the most enriched in adjacent tissue at the right. A students T-test was used to determine significance of the observed change in percentage: Cysteine cysteinylation is significantly enriched in the tumor (***p = 0.00045) while histidine oxidation (*p = 0.044), arginine citrullination (*p = 0.013), lysine ubiquitination (**p = 0.0031) and cysteine carbamidomethylation (**p = 0.0078) are significantly enriched in the normal tissue. (**B**) The percentage of overlapping sites between a randomly chosen subset of the cohort (30 peptides from 6 samples) and the remaining samples is shown. This was repeated 10,000 times to generate the intra-replicate distribution (light green; the mean is depicted as a dark green dashed line). The overlap for the identified phosphosites in the immunopeptidomics data and the CPTAC data is shown as a black line (top left). The abundance in the CPTAC cohort for the 5 overlapping phosphosites are shown in the tumor and adjacent tissue samples (wilcox p values for tumor vs. adjacent abundance is indicated in the figures). (**C**) Typically, antigenic peptides are classified by their genetic origin, including mutations, cancer-germline genes expressed outside of their biological context, oncogenic virus genes, genes with highly tissue specific expression patterns, or overexpression of genes with low endogenous expression (left block). In all these cases, PTMs can increase both the identification and therapeutic potential of these antigenic peptides. Likewise, PTMs can themselves be a source of antigenicity when pathways are activated in a disease-specific manner (right block) creating a PTM-driven antigenic epitope.

Although the frequency of the phosphorylation did not exhibit any significant differences between the tumor tissue and the adjacent controls, we found 27 phosphorylated peptides which only appeared in the tumor tissue and not in the adjacent control. We hypothesized that these tumor-specific phosphopeptides might originate from proteins that are phosphorylated more in breast tumor tissue. To examine this, we compared the immunopeptidomics to clinical phosphoproteomics data. Surprisingly, we could find that from the sites identified both in the immunopeptidomics and phosphoproteomics 42% were phosphorylated in both (Figure 6B). This is despite the fact that, on average, when comparing between different samples in the phosphoproteomics there is only a 37% overlap in phosphosites (Figure 6B). Furthermore, of the phosphosites identified in both cohorts, all were increased in the tumor compared to adjacent tissue, both on the phosphoprotein and HLA I-bound peptide levels (Figure 6B). This suggests that tumor-induced alterations of modifications on cellular proteins can propagate to changes in the presented landscape.

## Discussion

By developing PROMISE, we systematically analyzed the PTM landscape in the immunopeptidome and identified thousands of modified peptides across different cancers. Although numerous studies have examined HLA I presentation of modified peptides in the context of tumor antigenicity and autoimmune disease^6–10,61^, such analysis relied on experimental enrichment of the modification of interest. The capability to search large number of PTMs allowed us to identify types of modifications that were not examined before in the context of antigen presentation. For example, recent studies have suggested that the proximal ubiquitin may undergo proteasome degradation with its substrate^46,62^. Indeed, we could detect some remnants of ubiquitin-like modifications in our analyses, suggesting that they may be loaded and presented on MHC I and may serve as bone-fide antigens.

Modified-peptide analysis, coupled with structural modeling and binding assays, strongly suggests that modifications may generate novel HLA I binding motifs that could not be identified merely by the amino acid sequence. For example, cysteine is under-represented in HLA I ligand datasets^41^ hampering accurate binding predictions of cysteine-containing peptides^63^. However, by including several cysteine modification types in our search space, we could identify presented peptides containing cysteine with distinct motifs. Another example is the under-representation of unmodified lysine residues in the 2nd position anchoring site in the reported epitopes in the IEDB compared to the presence of modified lysine at this position. Notably, some of the peptides that we have identified do not match the consensus binding motif (8-11 mers). This may be driven by the PTM or reflect previous observations that have described longer peptide binding^64–70^. As binding motifs are the dominant selection criteria for antigen prediction algorithms^63,71^, PTM-driven motifs should prove invaluable to the next generation of binding prediction software^61^. It will be intriguing to examine PROMISE in the context of additional modifications. This type of analysis may also be extended to include mutations and the MHCII repertoire and non-canonical peptides such as ones generated by splicing, fusions or from non-coding regions ^28–32^.

Beyond expanding the HLA landscape, modified peptides may also signal changes in metabolic and signaling states of the cells under physiological or pathological circumstances. Notably, by comparing cancerous tissues and adjacent controls in TNBC, we found changes in the frequency of different modification types in the imunopeptidome. Further, we could confirm that 40% of the phosphorylation sites identified on HLA I-bound peptides also exhibited increased abundance in phosphoproteomics of breast cancer (CPTAC data). We note that the detection of the peptide may be due to both higher abundance of the protein or increased phosphorylation. Nevertheless, these results suggest that intracellular changes in the phospho-state of proteins, prior to degradation, may be kept and loaded onto HLA for presentation to create unique HLA signatures in breast tumors (Figure 6C). Although beyond the scope of this study, it also raises the intriguing possibility that drug-induced alterations in the activation of specific pathways may, in turn, alter the HLA repertoire. In the future, this feature may be utilized to direct immune responses against specific antigenic peptides in combination with targeted therapies ^1–4^.

Previous studies^6–12,72,73^ together with our analyses, highlight the potential of modified antigens to play an immunomodulatory role in tumor-host interactions and may drive either immune suppression or immune evasion. To further substantiate our predictions, we performed *ex vivo* and *in vivo* reactivity assays with several modified peptides and confirmed that they can elicit T cell cytotoxicity. While this class of modified antigens may offer novel therapeutic opportunities, there are important questions that remain to be addressed before these may be utilized for cancer therapy. For example, the tissue specificity of a potential immunogenic modification, the heterogeneity and stability of an altered modification state will need to be examined in the context of T cell recognition. Nevertheless, our analyses identified hundreds of modified testis antigens and tumor-associated peptides, which may serve as a new source of modified neoantigens in the context of immunotherapy. Coupled with patient-specific modifications, which occur sporadically and can be targeted for individualized therapy, we foresee that a broad range of potentially therapeutic antigens may be detected when analyzing peptide modification states. Beyond cancer, our approach may be utilized to expand our understanding of the PTM-driven HLA repertoire across different human pathologies, ranging from infectious diseases to autoimmunity and neurodegeneration.

## Material and Methods

### PROMISE

Current proteomics software focuses on data from samples where an exogenous enzyme, like trypsin, was used to digest the proteins into peptides. This reduces the potential search space to only peptides with either lysine (K) or arginine (R) terminal residues. By contrast, HLA class I peptides are cleaved by the proteasome and a number of endopeptidases, generating peptides that are between 8 and 15 amino acid residues and may have any terminal residue. Computationally, this means that the search space for endogenously-cleaved peptides with modifications must contain every potential protein fragment with multiple potential mass shifts, leading to an exponential growth of the search space and making the duration of the search challenging^74^. PROMISE (PROtein Modification Integrated Search Engine) optimizes search efficiency with two stages: a) matching phase and b) prioritizing phase (supplementary pipeline documentation). The matching phase reduces the algorithm running time, utilizing the ultrafast MSFragger^33^ software and parallel computing on a CPU cluster. The prioritizing phase includes several computational steps to distinguish between true and false hits, validate PTM identifications and site position and rank predictions by their biological relevance and antigenic potential. To evaluate pipeline performance, we used the full human proteome from UniProtKB as reference data and searched for endogenous proteasome-cleaved peptides^75^ (length between 6 and 40 amino acids) with 5 variable modifications, creating a search space of ~31 billion potential peptides. To assess the reproducibility of the identified peptides by the distributed version and the standalone MSFragger we compared the spectral assignments from identical sets of data and found that 99.2% were identical. We then compared the identifications in PROMISE to those with the original search criteria (standard search: only n-acetylation and methionine oxidation included). In cases where the peptide to spectrum matches (PSMs) conflicted between the two searches (1.22% of PSMs),PROMISE prioritizes the highest scoring match. Although the scoring alone is not a guarantee of a true assignment, it does suggest that the inclusion of a modification in the predicted peptide better described the spectrum.

For the paper analysis we used a subgroup FDR whereby we split the identifications into three groups: unmodified, standard search modification types (n-acetylation and methionine oxidation) and the other modification types. For the MC38 immunopeptidomics where the cohort was too small to successfully execute subgroup FDR (Figure 4) and where additional enrichment analysis where being performed (Figures 2, 3 and 6A) we used a global FDR. In both cases, the cutoff was set to 5%. In cases where subgroup FDR was used across multiple cohorts, we included any peptide that passed the subgroup FDR in at least one cohort. Detailed software architecture and performance can be found in the supplementary pipeline documentation.

### Modification Annotation and Classification

In order to assess the effects of modifications in a holistic manner, we considered both modifications that may arise during sample processing or handling and ones that reflect an altered cellular state (“biological”). This was done using the UNIMOD classification system (unimod.org). We explicitly note that the nature of the modification is not sufficient to determine whether the modified form was generated due to biological regulation or whether the peptide is presented on its modified form. As peptides may exist in the cell in either their modified or unmodified form, we chose for validation only peptides that were significantly different between the cancerous and control conditions. When a peptide contains multiple modification types, we defined a leading modification, prioritizing ‘biological’ modifications over some that may be considered as technical (based on the unimod classification).

### Search mass boundary effect correction

The search space in the analysis is bounded by a 15 amino acid peptide length. This can result in incorrect assignments when a contaminant with a mass higher than 15 AA is assigned to a 15-mer peptide with a high mass shift modification. As we search for PTMs with large mass shifts (e.g. ubiquitin tail with 4 amino acid GGRL - 383.228103 Da), this can lead to mis-assigned spectra. Because the longer peptide is not part of our search space we cannot rule out that a better match exists or that there is a higher scoring match above 15 AA. Therefore, to avoid a bias we filter out potential mis-assignments by limiting the total peptide mass to the average mass of 15 amino acid peptide plus 100 Da when comparing peptide lengths (Figure 1G).

### HLA I motif

HLA I motif presentation was designed to capture both the main anchor position 2 and C-terminus and the TCR recognition area (position 3-7). The presented motif was created by collecting all the epitopes reported for the specific HLA haplotype from the IEDB^42^ database. Epitopes with length less than 8 amino acids were discarded. To correct for discrepancies in length, the motif was constructed from positions 1 to 7 starting from the N terminus followed by the C terminus and its preceding position. For 9 mer epitopes, the motif is taken from all 9 positions, for 8-mer epitopes the 7^th^ position is duplicated and presented as both positions 7 and 8/C-1. For epitopes longer than 9 residues, the motif skips positions 8 till C-terminus −1. Motif logos were plotted using Seq2Logo 2.0^76^ with default parameters. The comparable motif was created using Two-Sample-Logo^77^

### Site score

The score was designed to determine if a PTM tends to fall within the peptide anchor positions or the center positions (3-7) of the peptide. By summing up the differences between the distribution values of modified amino acids vs the background in the anchor positions (2, C-terminus) and subtracting the sum of distribution differences in the center positions (3-7). An enrichment in the anchor positions will result in a high positive score while enrichment in the center of the peptide will result in a negative score. In case both the center and anchor positions are enriched or under-represented, the score will be close to zero and we cannot classify the modification tendency to be in a specific area.

### TCGA, CPTAC Phoshoproteomics, and Immunopeptidomics Analysis

The cancer genome atlas (TCGA) data was mined using the xenaPython package in Python 3.6. The results shown in this analysis are in whole or part based upon data generated by the TCGA Research Network: http://cancergenome.nih.gov/. Colon Adenocarcinoma (COAD), Breast Cancer (BRCA) Skin Cutaneous Melanoma (SKCM), and Glioblastoma (GBM) cohorts data were used. Data used in this publication were generated by the Clinical Proteomic Tumor Analysis Consortium (NCI/NIH). The CPTAC Breast Cancer phosphoproteomics data ^14^ was compared to the Triple-Negative Breast Cancer immunopeptidomics data ^40^. The CPTAC intra-replicate site overlap was calculated from the tumor samples in the cohort by randomly drawing 30 phosphosites from 6 samples in the same TMT experiment and comparing the identification to the remaining TMT experiments. This was done 10,000 times and is presented in Figure 6B. The overlap between the CPTAC phosphoproteomics and immunopeptidomics was defined as the number of phosphosites identified in both CPTAC and immunopeptidomics data (n = 5) out of the sites which were covered by peptides in both datasets (n = 12). The remaining 18 sites only had peptides covering them in the immunopeptidomics data and therefore could not be evaluated in the tryptic CPTAC cohort.

### Modeling the peptide-receptor complex

General modeling scheme –The FlexPepBind scheme used here^78,79^ allows the structure-based evaluation of the relative binding affinities of different peptides for a given receptor, using a solved structure of a representative peptide-protein interaction as a template. Structures of peptide-MHC complexes were generated by “threading” candidate peptide sequences onto this template, followed by refinement using Rosetta FlexPepDock^52^. The top-scoring models were selected to discriminate stronger from weaker binders and inspected for the structural details of an interaction.

#### Selection of templates for modeling

For each of the MHC alleles (receptors) and peptides, we evaluated different available PDB structures to serve as templates for the modeling of the structure and relative binding affinities of different peptides. Screening for relevant PDB templates was guided by 3 main requirements: (1) matching MHC allele, (2) matching peptide length, and (3) similarity of peptide anchor residues. Specifically, for peptide K(ac)P(ox)SLEQSPAVL bound to HLA-A02 (Figure 3H) we used PDB id 5D9S^80^ (HLA-A02 bound to FVLELEPEWTV); for peptide KP(ox)LKVIFV bound to HLA-A02 (Supp. Fig 1), we used the peptide backbone from PDB id 4F7T^81^ (HLA-A24 bound to RYGFVANF) and the same MHC receptor structure (from PDB id 5D9S); for peptide MPTLPPYQ(me) bound to HLA-B54 (Figure 3I), we used PDB id 3BWA^82^ (HLA-B35 bound to FPTKDVAL). Residues that differ between the MHC alleles were “mutated” using the fix backbone protocol (Rosetta fix_bb; [8]); for peptide TLIESK(me)LPV bound to HLA-A02 (Figure 5C), we used PDB id 3MRK (HLA-A02 bound to PLFQVPEPV).

#### Modeling peptide onto MHC receptor using the selected template

Using the Rosetta fixbb protocol for fixed backbone design^83^, we modeled the desired peptide sequence onto the template peptide, while keeping the side chains of the receptor fixed. We then used Rosetta FlexPepDock refinement in full-atom mode to optimize the structure of the complex with the threaded target peptide (all peptide atoms, as well as the receptor interface sidechains, were allowed to move). For each sequence, we generated 200 models. These were scored, and the 5 top-models were selected to represent the MHC-peptide interaction of interest. Comparison of the top scoring models of the modified peptides and corresponding non-modified peptides allowed inspection of the atomic details of their differential binding.

#### Scoring function

The standard Rosetta score function^84,85^ was used, and models were assessed according to their FlexPepDock reweighted score (sum of Total score, Interface score and Peptide score; where Total score is the overall Rosetta energy score for the complex, Interface score is the energy of pair-wise interactions across the peptide-protein interface and Peptide score is the sum of the Rosetta energy function over the peptide residues). This score was shown to discriminate well near-native structures in previous FlexPepDock modeling studies^86^.

### ProImmune binding assay

ProImmune (https://www.proimmune.com) Module 2 REVEAL Binding Assay measures the yield of correctly conformed MHC-peptide complexes following incubation of the recombinant MHC allele with and the peptide of interest, using a conformation-dependent antibody in an immunoassay. Each peptide is given a score relative to the positive control peptide, which is a known T cell epitope.

### Reagents

A complete list of reagents, antibodies & Chemicals can be found in supplementary data 6.

### Purification and analysis of the MHC peptides from MC38 cells

MC38 cells were kindly provided by Ayelet Erez (Weizmann Institute). H2-Kb and H2-Db -bound peptides were isolated from three independent preparations of MC-38 cell line, each containing 5e8 cells, as in (Milner et al. 2013). Briefly, cells were lysed with lysis buffer comprised of PBS supplemented with 0.25% sodium deoxycholate, 0.2 mM iodoacetamide, 1 mM EDTA, 1:200 protease inhibitors cocktail (Sigma, St. Louis, MO), 1 mM PMSF, and 1% octyl-β-D-glucopyranoside. The lysate was then shaken on a shaking table gently for one hour at 4°C, cleared by centrifugation at 4°C and 47,580g, for 60 min (Sorval RC 6+ centrifuge, Thermo Fisher Scientific). After centrifugation, the supernatant was passed through a column containing the Y3 antibody (anti-H2-Kb) or 28-14-8 antibody (anti-H2-Db) covalently bound to protein G Sepharose resin with dimethylpimelimidate. Next, the columns were preconditioned with two column volumes of 0.1 N acetic acid, and next with two column volumes of 20 mM Tris/HCl, pH 8.0. After passing the cleared cell extracts, the columns were washed with five column volumes of 400 mM NaCl and 20 mM Tris-HCl pH 8, followed by another wash with 20 mM Tris-HCl, pH 8. The MHC-bound peptides were eluted with 1% trifluoroacetic acid, desalted, concentrated and separated from the MHC molecules by reversed-phase fractionation using disposable Micro-Tip Columns C-18 (Harvard Apparatus, Holliston, MA). The peptides were eluted with 30% acetonitrile in 0.1% TFA, dried by vacuum centrifugation, and dissolved in 0.1% TFA for analysis by capillary chromatography combined with tandem mass spectrometry (LC-MS/MS). Samples were resolved by capillary chromatography using an UltiMate 3000 RSLC coupled by electrospray, to a Q-Exactive-Plus mass spectrometer (Thermo Fisher Scientific). Elution of the peptides was performed with a linear two-hour, 5-28% acetonitrile gradient in 0.1% formic acid, at a flow rate of 0.15 μl/min. The 10 most intense ions in each full-MS spectrum, with single to triple-charged states, were selected for fragmentation by higher energy collision dissociation (HCD), at a relative collision energy of 25. Ion times were set to 100 msec. automatic gain control (AGC) target was set to 3*106 for the full MS, and to 1*105 for ms2. The intensity threshold was set at 1*104.

### MS Spectra validation and visualization

Modified peptides were synthesized through Peptide 2.0 company in purification level above 95%, then synthesized peptides were analyzed in mass spectrometry using target search mode. For asparagine deamidation we synthesized the modification as aspartic acid. The spectrum comparison visualization and a similarity score between the original spectrum and the synthesized spectrum are created by R package OrgMassSpecR. Thermo Xcalibur Qual Browser was used to manually annotate spectra. Spectra visualization is created using PDV 1.5.4 software^87^ including a,y,b ions and all potential losses.

### *Ex vivo* and *In vivo* peptide specific killing assays

Animal experiments were approved by the Institutional Animal Care and Use Committee (IACUC) of the Weizmann Institute of Science (approval number 05250620-2).

#### BMDC generation and immunization

Bone marrow derived dendritic cells were generated according to ^56^. In brief, tibia and femur of donor mice were isolated, and the bone marrow flushed out into PBS using a syringe. Clusters were dispersed by pipetting, and cells were washed in PBS and seeded in 10 mL RPMI, 10% FCS (heat inactivated), with 1 mM sodium pyruvate, 2 mM glutamine, 1x non-essential amino acids, 1 mM HEPEs, and 50 uM beta-mercaptoethanol, 200U/mL GM-CSF (Peprotech) onto petri dishes (day 0). At day 3, 10 mL fresh medium was added. On day 6, 15 mL of the medium was removed, floating cells collected by centrifugation (200x*g*, room temp, 10 minutes), resuspended in 5 mL and added back to the plates with fresh GM-CSF. On day 8, loosely attached and floating cells were collected and reseeded at 10-15e6 cells per tissue culture plate with 100 U/mL GM-CSF. Cells were matured by addition of LPS (LPS, 1 ug/mL) on day 9 and harvested by day 10. Successful differentiation was confirmed by staining cultures with CD11c, MHC II, CD80 and CD86.

Floating cells were collected, washed in PBS and resuspended in OptiMEM (Gibco) to 10e6 cells/mL. Peptides (3x unmodified, 3x modified counterparts) were added at 200 uM, and the cells incubated at 37C for 3-5 hours for loading. DCs were finally washed, resuspended and pooled to either unmodified immunized or modified immunized. Cells were injected to female C57Bl/6J mice i.p. in 200 uL PBS. Immunizations were repeated weekly for a total of 3 times.

#### Ex vivo tumor killing assay

Lymphocyte sensitization and cytotoxicity assays were adapted and modified from^88–91^. Specificly, 1e5 MC38 cells were injected to mice one to two weeks after the last of three immunizations with either unmodified or modified peptides, or to unimmunized mice as control. After 19 days, spleens were collected and single cell suspensions were generated and pooled together from four mice per each group. To sensitize peptide-specific T cells, the pooled splenocytes were split into separate groups for each specific peptide, and 1/3 of the total cells in each group was pulsed with 50 ug/mL peptide for 2 hours at 37C, and added to the remaining 2/3 of the splenocytes. Pulsed cells were cultured for 5 days (termed sensitized lymphocytes). Splenocytes from non-immunized tumor-bearing mice (non-immunized control) were cultured without addition of peptides for the same period (termed ‘control lymphocytes’).

One day before the assay, MC38 cells were stained with CFSE (5 uM) for target cell identification and seeded at 1e4 cells/well in a 96 well plate. Sensitized or control lymphocytes were collected and enriched using Lympholyte M (Cedarlane Labs) according to the manufacturers protocol. Finally, lymphocytes were counted and seeded onto CFSE-labeled MC38 target cells at different Target:Effector ratios (1:12.5, 1:25, 1:50).

After 18 hours, supernatants were transferred, adherent cells collected with trypsin and unified with the supernatant containing dead cells. The cell suspension was diluted with PBS containing Propidium iodide (final concentration1 ug/mL) and read on an Attune flow cytometer (Thermo Fisher).

Cells were gated for single cells, then target cells identified by CFSE staining. Dead cells were discriminated by PI positive and negative (live) gating (see Supp. Fig 6A). We determined before co-culturing the amount of CD8 cells in the cultures by flow cytometry (non-immunization control 31.9%, unmodified immunized 47.9%, 45%, 46.3%, modified immunized 47.4%, 42.6%, 45.9%). The killing rates of non-immunized mice were corrected for the difference in CD8. The specific killing was calculated by the formula (as described in ^90^)

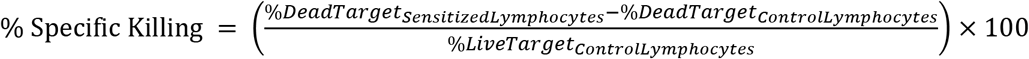

#### In vivo peptide-specific killing assay

Mice were immunized by DC-loaded peptides as described above and one to two weeks after the last of three immunizations, 1e5 MC38 cells were injected to boost the immune response.

Target cell preparation: Single cells suspensions from naïve CD45.1 spleens were pooled and pulsed with 50 ug/mL peptides (unmodified, modified, or left unpulsed) for 3 hours at 37°C. Cells were washed twice and each experimental group (ummodified, modified, unpulsed) was differentially stained with either CFSE (Biolegend), CellTracker Deep Red (Thermo Fisher Scientific), or Tag-it Violet (Biolegend) cell tracking dyes for 25 minutes. Three different concentrations, high, medium, and low) of each dye were examined (CFSE 5 uM, 0.5 uM, 0.05 uM; CellTracker Deep Red 4 uM, 0.4 uM, 0.04 uM; Tag-it Violet 10 uM, 0.1 uM, 0.01 uM). Staining reactions were quenched with PBS/FCS 7.5%, washed twice and counted. Cells in each dye group (high, medium, low) were mixed at 1:1:1 ratios for a mix of differentially labeled unmodified-pulsed, modified-pulsed, or unpulsed splenocytes), and 45e6 cells per mouse were injected i.v. in 200ul PBS into MC38 tumor-bearing mice. Two naïve mice were injected with the same peptide loaded splenocytes as controls.

After 18 hours, the spleens were harvested, and processed for single cell suspensions and stained with CD45.1 PE-Cy7 (Thermo Fisher Scientific) after incubation with an Fc blocking antibody (Biolegend). Finally, cells were washed, resuspended in PBS and fixed with Formaldehyde (end concentration 1%). Samples were acquired on an Attune NxT flow cytometer and analyzed with Flowjo.

Cells were gated for lymphocytes, singles cells and CD45.1 (see gating strategy Supp. Fig 4A). Target populations were identified by CFSE, CellTracker Deep Red or Tag-it Violet signal, then each peak gated to determine their percentages. In vivo killing was calculated according to ^55,56^.

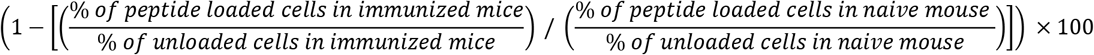

### MSFragger search parameters

Search params were set to default for close search with the following changes: Precursor true tolerance was set to 10 ppm; fragment mass tolerance was set to 20 ppm. Search enzyme was set to nonspecific enzyme with cleavage after ARNDCQEGHILKMFPSTWYV. Peptide lengths were set between 8 and 15. Num enzyme termini = 0, clip nTerm M = 1, allow multiple variable mods on residue = 0, max variable mods per mod = 3, max variable mods combinations = 65000.

### Bioinformatics and data analysis

Statistical analyses were performed in Prism 8 software (GraphPad, San Diego, CA, USA) and R v 3.6.1. heatmap was drawn with pheatmap 1.0.12 and ComplexHeatmap 2.2.0 R package with Euclidean distances for clustering where relevant. Flow cytometry data were analyzed with FlowJo V10 from Becton, Dickinson, and Company. Experimental schematics were generated using BioRender.

## Data and code availability

MC38 immunopeptidomics data was deposit in PRIDE archive with ID PXD017448 and standard MaxQuant^92^ analysis results. All the code used is available from the corresponding author upon reasonable request.

## Acknowledgment

We thank the members of the Merbl lab as well as Cyrille Cohen and Michal Besser for scientific discussion; Irun Cohen, Avital Eisenberg-Lerner, Julie DeMartino, Chaim Putterman and Einat Zisman for critical reading of the manuscript. We would like to thank Sara Meril for technical help and Ayelet Erez for MC38 cells. Y.M. is supported by the European Research Council (ERC) under the European Union’s Horizon 2020 research and innovation program (grant agreement No 677748); The I-CORE Program of the Planning and Budgeting Committee and The Israel Science Foundation (Grant No. 1775/12) and the Israeli Science Foundation (Grant No. 2109/18); The Gruber Peter & Patricia award. The US national Institutes of Healths grants R01-GM-094231 and U24-CA210967 (to A.I.N). Y.M. the incumbent of the Leonard and Carol Berall Career Development Chair. This research was partially supported by the Israeli Council for Higher Education (CHE) via the Weizmann Data Science Research Center and by a research grant from Madame Olga Klein – Astrachan.

## Author contribution

A.K. and A.J. led the study and performed all computational analyses unless otherwise mentioned. M.P.K and A.S. performed in vitro and in vivo assays and their analysis. T.S. performed 3D modeling, D.M. and Y.L. consulted regarding Mass Spectrometry analyses and algorithm development. E.B. generated the HLA I peptidomics data and A.K., G.C.T F.V.L and F.Y performed software development. G.C consulted regarding validation assays O.S.F., A.A., L.E. and A.I.N supervised the work of respective group members and A.J., A.K. and Y.M. wrote the manuscript and Y.M. supervised the study.

