## Supplementary data 1 for "Uncovering the modified immunopeptidome reveals insights into principles of PTM-driven antigenicity"

**Table 1. Description of the different combinations of PTM types and amino acids comprising the set searched against in our analysis:**

| Modification name | Amino acid | Mass shift | UNIMOD Accession # | UNIMOD classification | Remark |
|---|---|---|---|---|---|
| Methionine oxidation | M | 15.99490 | 35 | Artefact | Common chemical non-enzymatic modification. Appears in most MS searches [95] |
| protein N-termini acetylation | [X]@N-terminus | 42.01060 | 1 | Multiple | |
| Phosphorylation | YTS | 79.96633 | 21 | PTM | |
| Acetylation | K | 42.01060 | 1 | Multiple | [96] |
| Methylation | CHNQKRILDE | 14.01565 | 34 | PTM | |
| Di-methylation | KR | 28.0312 | 36 | PTM | |
| Oxidation | WHKPC | 15.99490 | 35 | KPC – PTM<br>WH - Artefact | [97] |
| Deamidation | NQ | 0.98402 | 7 | Artefact | NQ - Artefact |
| Citrullination | R | 0.98402 | 7 | PTM | Enzymatic modification |
| Ubiquitination | K | 57.0215 (G)<br>114.0429 (GG)<br>270.144 (GGR)<br>383.228103(GGRL) | 1263<br>121<br><br>535 | Other<br>Other<br><br>Chemical derivative | |
| Sumoylation | K | 215.0906 (GGT)<br>343.149184 (GGTQ) | <br>1293 | <br>Other | G and GG – cannot distinguish between ubiquitin, Sumo or FAT10 |
| FAT10 | K | 227.127 (GGI)<br>330.136176 (GGIC) | <br>1990 | <br>PTM | |
| Cysteinylation | C | 119.004099 | 312 | Multiple | |
| Carbamidomethyl | C | 57.021464 | 4 | Chemical derivative | Artefact – used as fix modification in trypsin digestion |
