## Supplementary figures 1-6 for "Uncovering the modified immunopeptidome reveals insights into principles of PTM-driven antigenicity"

A

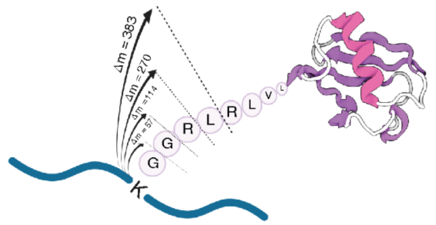


B

| Modification name | UNIMOD Accession # | Mass shift | Amino acid | UNIMOD classification | remark |
|---|---|---|---|---|---|
| Ubiquitination | 1263<br>121<br><br>535 | 57.0215 (G)<br>114.0429 (GG)<br>270.144 (GGR)<br>383.228103(GGRL) | K | Other<br>Other<br><br>Chemical derivative | |
| Sumoylation | 1293 | 215.0906 (GGT)<br>343.149184 (GGTQ) | K | Other | G and GG – cannot distinguish between ubiquitin, Sumo or FAT10 |
| FAT10 | 1990 | 227.127 (GGI)<br>330.136176 (GGIC) | K | PTM | |

C

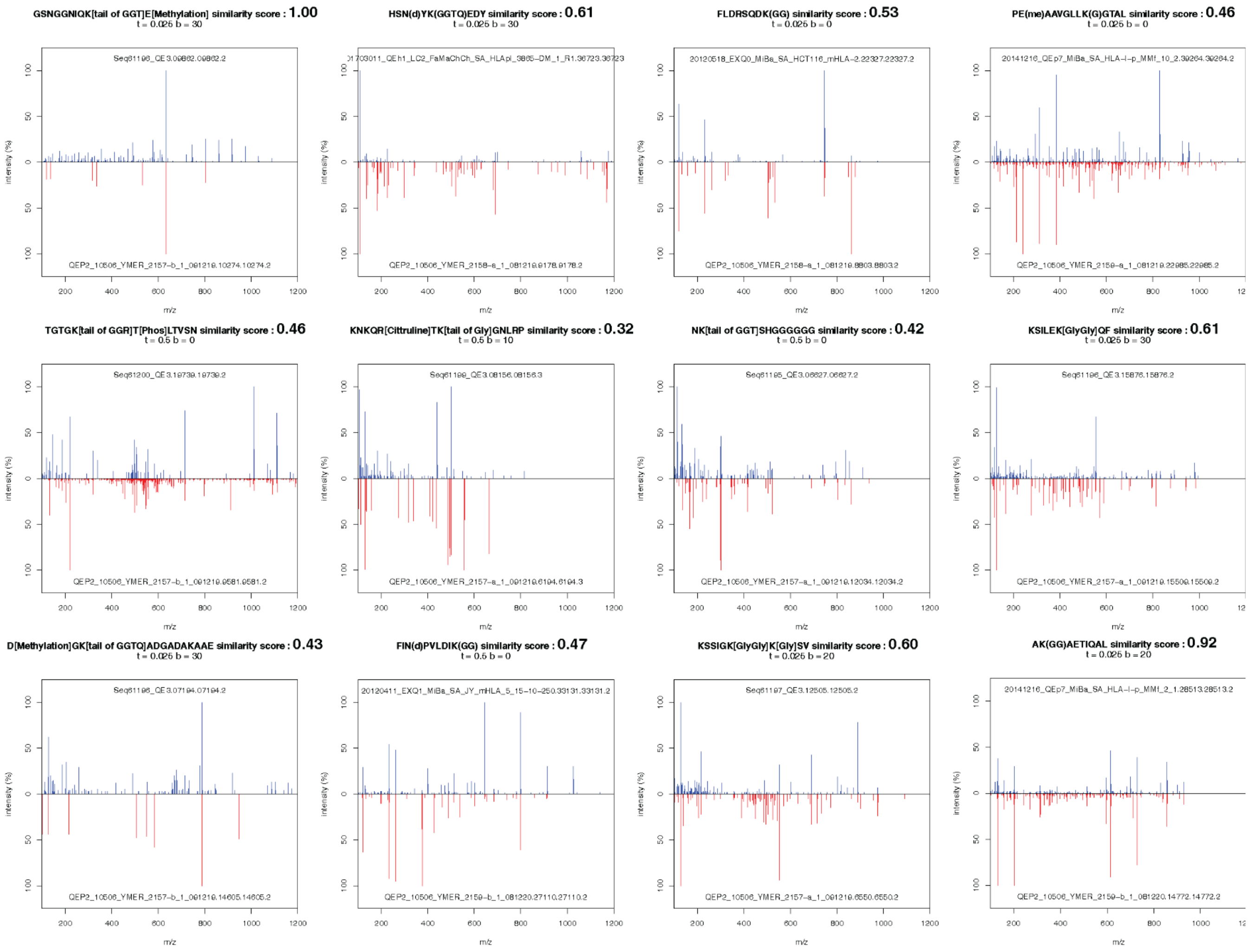


**Sup – Figure 1. Ubiquitin tail search on immunopeptidomics data**

(**A**) The search of ubiquitin tail on endogenous HLA peptides requires a definition of any tail length as a variable mass shift. (**B**) List of different ubiquitin family tails in the multi-modification search (**C**) examples of modified peptides with a different type of ubiquitin tails identified in the datasets, each peptide was synthesized (Peptide 2.0 Inc) and its spectrum was capture using mass spectrometry. A similarity score was calculated between the synthesized spectrum (red) and the original spectrum in the dataset (blue) using R package OrgMassSpecR.

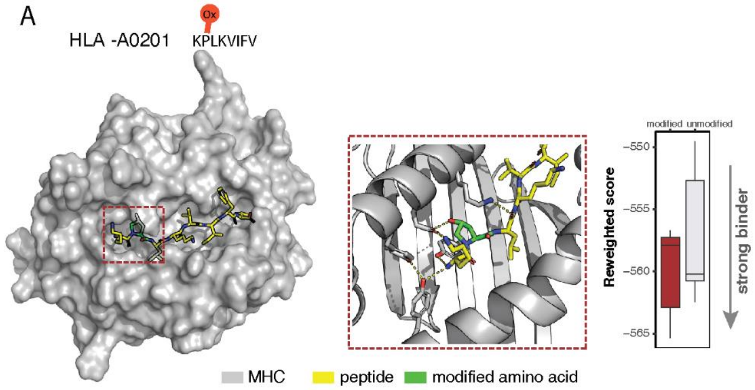


**Sup-Figure 2. KP(ox)LKVIFV and HLA-A0201 3D interaction**

(**A**) Rosetta FlexPepDock structural model of the interaction between the modified peptide KP(ox)LKVIFV (yellow sticks) and the MHC molecule haplotype HLA-A0201 (grey surface \ cartoon). The modified amino acid (green) creates a more stable interaction with the MHC molecule as compared to the unmodified form. The effect of the modified amino acid is shown in detail in the zoom-in picture. The proline hydroxyl group at position 2 forms a stabilizing hydrogen bond with MHC receptor residue E-87 (shown as dashed yellow line, as well as other hydrogen bonds between peptide and receptor). FlexPepDock reweighted score was calculated for the interaction between the MHC and modified or unmodified peptide. A more negative score indicates a more stable interaction

A

| peptide | Similarity score |
|---|---|
| GSNGGNIQK[tail of GGT]E[Methylation] | 1 |
| Ala(ac)-ATTGSGVKVPRNF | 0.999633 |
| YFGQG-R(me2)GGGGGGGY | 0.99531 |
| S-Arg(Me)-GGG-Asp(Me)-QGYGSGR | 0.929709 |
| SY-Arg(Me2)-GNYGGGGGG | 0.897688 |
| GFGVGR(me2)GITTP | 0.896315 |
| M(ox)DK(ac)NELVQKA | 0.759881 |
| GGSR(me)GFGGGGY | 0.548236 |
| Ala(ac)-SSDIQVKELEKR | 0.472082 |
| NK[tail of GGT]SHGGGGGG | 0.415029 |
| TAVACR[Citrullination]LTQM(ox) | 0.349643 |
| KSYS(p)FIARM | 0.311854 |
| GSG-pS-V-Arg(Me2)-GGGG | 0.220585 |
| TALFHTD[Metylation]D[Methylation]I | 0.131332 |
| SAVVAT[Phospho]VVVQ[Methyl]P | |
| RNNFGGDYR[dimety]GSSGD | |
| GAS[Phos]VS[Phos]ENHEIY | |

similarity score

similarity score
+ manual annotation

manual annotation

B

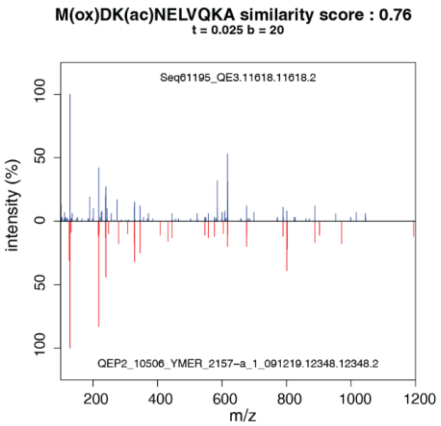


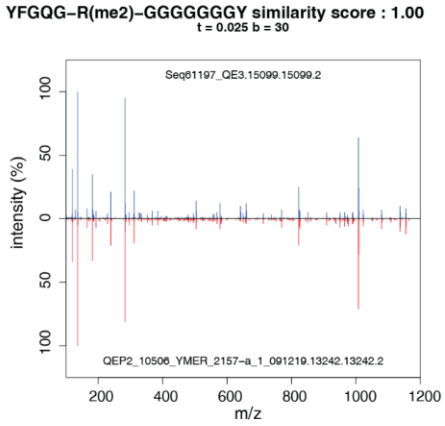


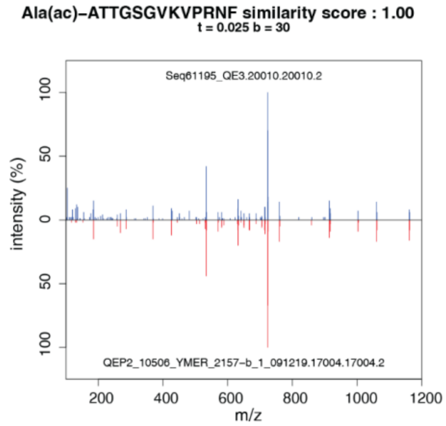


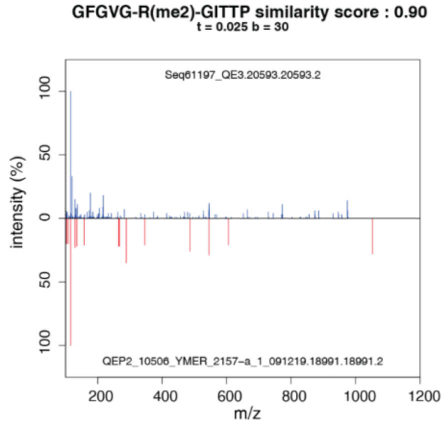


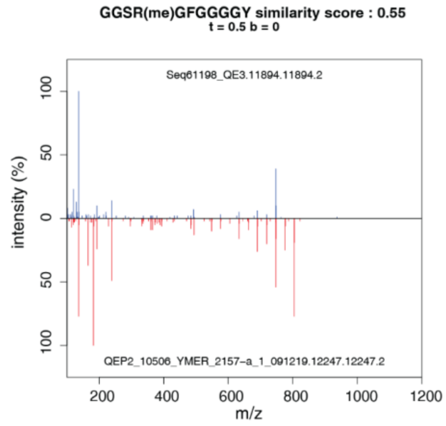

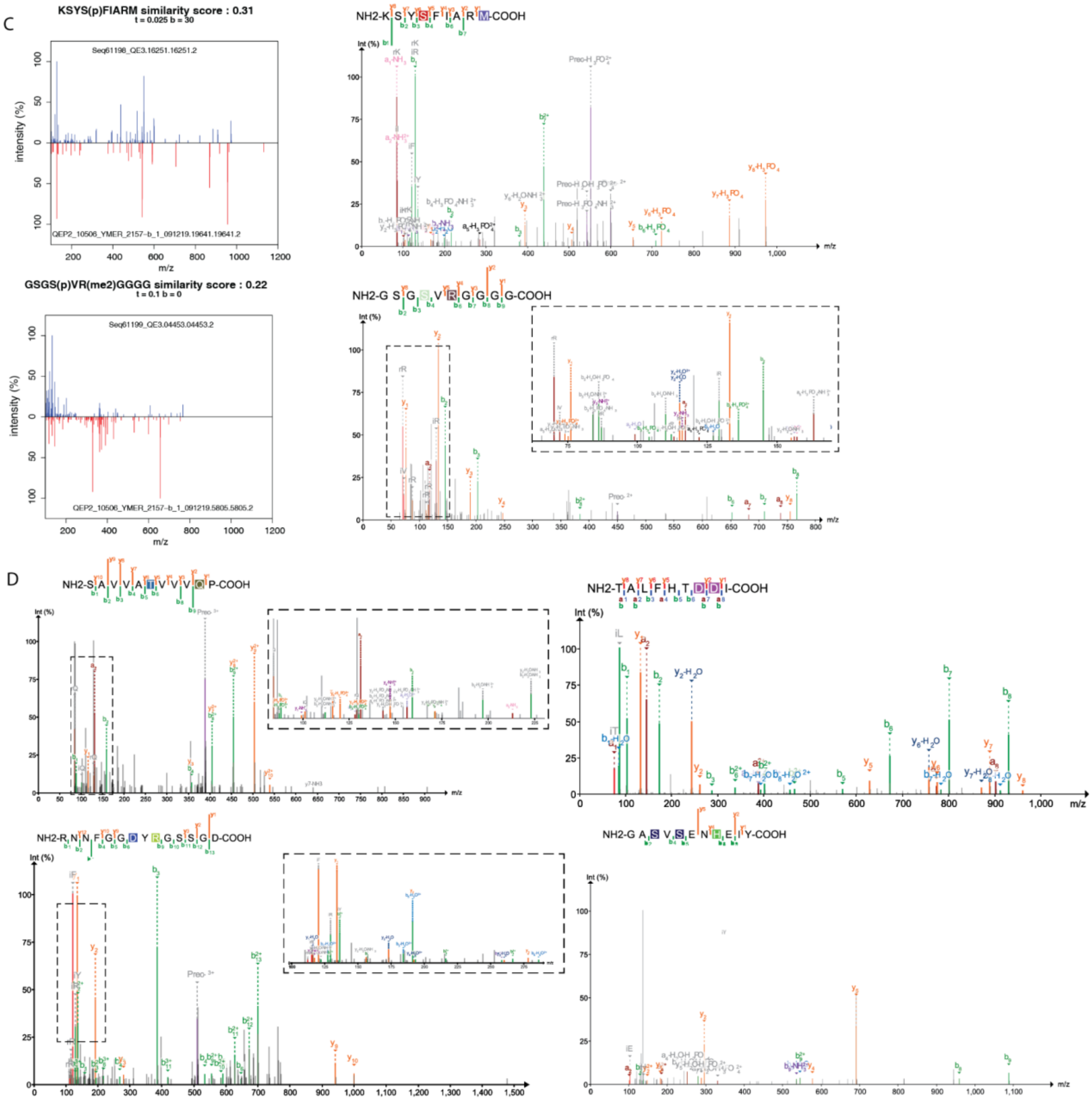


**Sup – Figure 3. Spectra validation**

(**A**) Modified HLA peptides, identified in MC38 cell line and not in healthy mouse colon tissue or reported in the IEDB dataset, were synthesized (Peptide 2.0 Inc) and their spectra were captured using mass spectrometry. For each modified peptide, a similarity score was calculated between the synthetic spectrum and the original spectrum using R package OrgMassSpecR. For a similarity score below 50%, manual annotation was done to validate the spectra. For peptides which we fail to produce a synthetic spectrum, we used manual annotation to validate. (**B-D**) spectrum comparison visualization and a similarity score are created by R package OrgMassSpecR, synthesized spectrum (red) in a mirror image of the original spectrum in the dataset (blue). In case manual annotation was done, visualization is created using PDV software[2] including a,y,b ions and all potential losses. (B) Validation based on similarity score (C) Validation based on both similarity score and manual annotation. (D) Validation based on manual annotation.

A

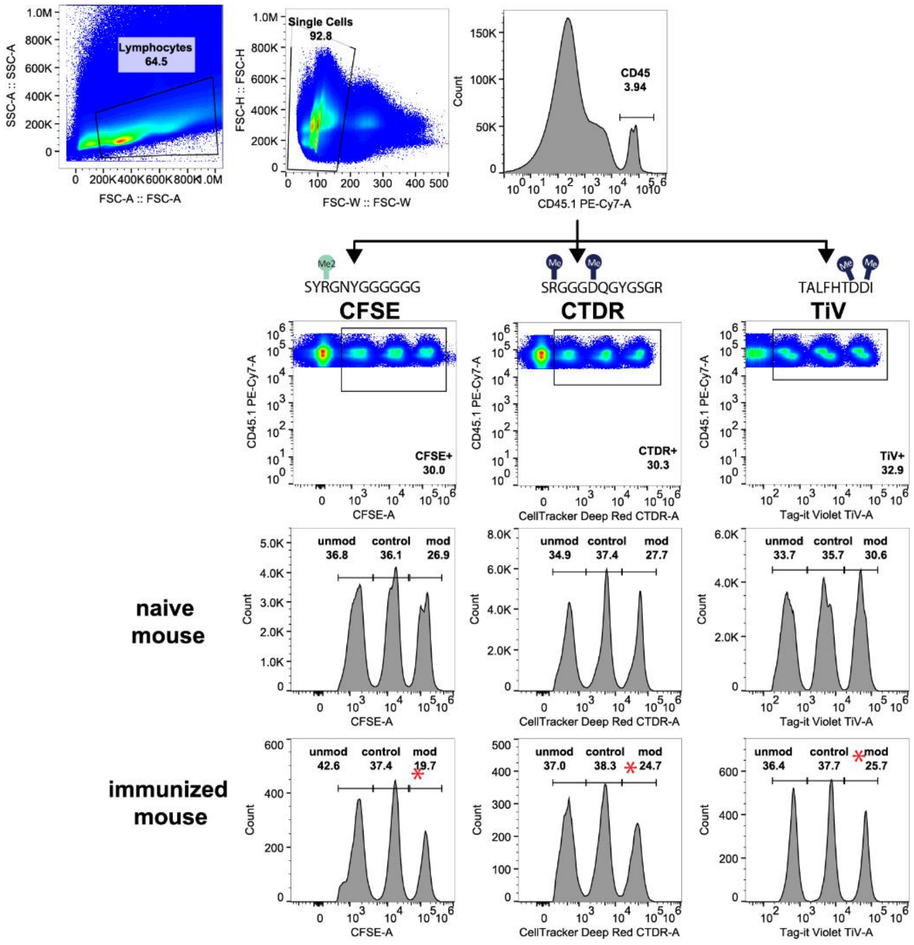


**Sup – Figure 4. Flow cytometry gating strategy for killing assays**

**(A)** The gating strategy to identify the 3 differentially labeled populations in the in vivo peptide-specific killing assay. The samples were stained with cell tracker dyes at low (non-modified peptide), high (modified peptide), and intermediate (un-pulsed) concentrations and injected to mice. Each non-modified/modified combination got one of three-tracker dyes (CFSE, CellTracker Deep Red (CTDR), or Tag-it Violet (TiV)). After 18 hours, the spleens were analyzed, and the percentage of killing was calculated normalized to naïve mice injected with the same peptide loaded spleenocytes as the immunized mice. Results from one representative immunized and naïve mouse are shown, the red asterisk indicates the peak corresponding to the modified-loaded splenocytes.

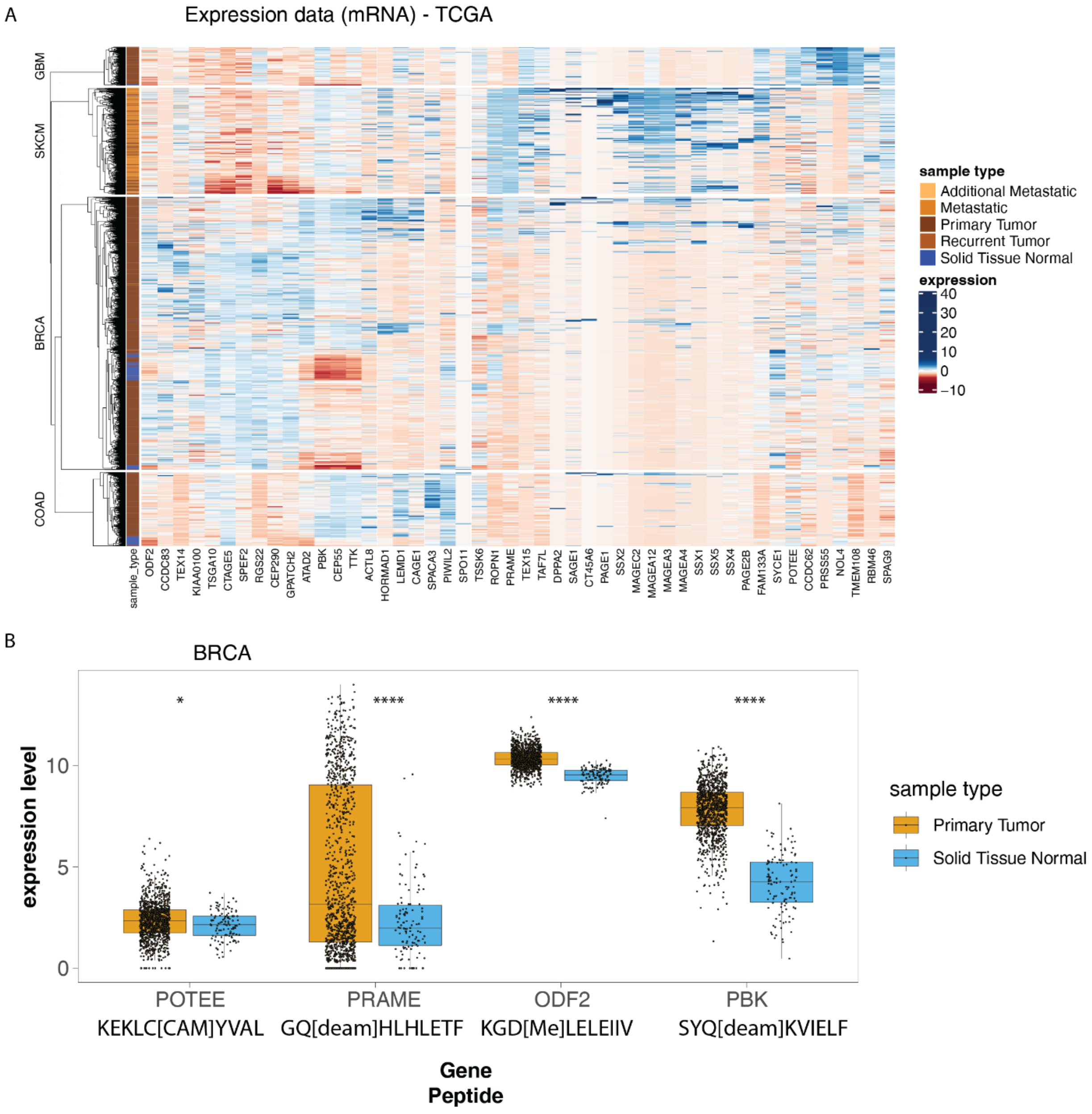


**Sup - Figure 5. Expression of genes encoding for testis antigens identified in PROMISE**

(**A**) TCGA mRNA expression data of testis genes in four different cancer type from which patient sample or cell lines immunopeptidomics data was analyzed by PROMISE: COAD – HCT116; BRCA – PXD009738, HCC1143 and HCC1937; SKCM – PXD004894; GBM- PXD003790. (**B**) The expression of the parent gene from 4 modified HLA I-bound peptide identified in PROMISE is shown for TCGA expression data from BRCA primary tumor and normal tissue. The parent testis gene is significantly overexpressed in the tumor tissue vs the normal (wilcox p values for tumor vs. adjacent abundance indicated in figures).

A

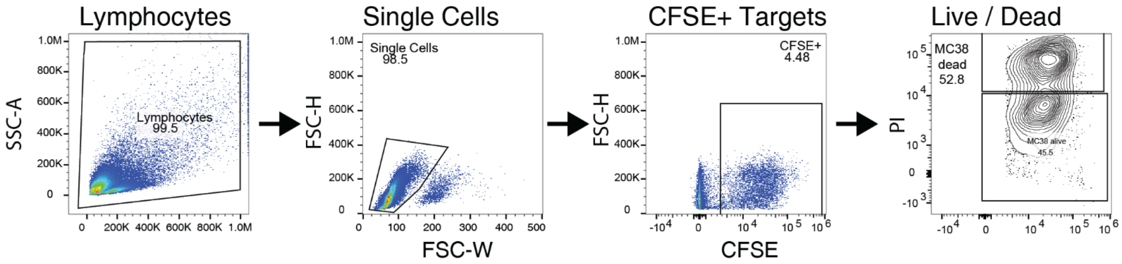


**Sup – Figure 6. Flow cytometry gating strategy for peptide ex vivo killing assay**

**(A)** Gating strategy for identification of dead target cells in the ex vivo tumor killing assay. All cells were included, only doublets were excluded from the analysis by FCS-W/FSC-H parameters. Dead cells were determined by CFSE positivity and PI positive staining.
